# The decidual microenvironment shapes neutrophil subsets that promote extravillous trophoblast invasion in early pregnancy

**DOI:** 10.64898/2026.11.16.766955

**Authors:** Zala Nikita Juvan, Oliver Kindler, Teresa Kloimböck, Kathrin Maitz, Nejra Cosis- Mujkanovic, Eva Achhammer, Anna-Lena Höbler, Dorien Feyaerts, Désirée Forstner, Sofia Raftopoulou, Ana Santiso, Sigrid Vondra, Andreas Ian Lackner, Gerit Moser, Julia Feichtinger, Martin Gauster, Christina Stern, Akos Heinemann, Jürgen Pollheimer, Christian Wadsack, Herbert Fluhr, Brice J. Gaudilliere, Julia Kargl

## Abstract

Successful establishment of the maternal-fetal interface is a critical process during early pregnancy, ensuring healthy fetal development. A critical phase in establishing the utero-placental blood supply is the invasion of extravillous trophoblasts (EVTs) into the maternal spiral arteries, marking the transformation of the endometrium into the decidua. By separating the EVT invaded decidua (iDec) and EVT non-invaded decidua (niDec), we demonstrate that neutrophils dominate the iDec immune landscape during early human pregnancy. High-dimensional flow cytometry and single-cell transcriptomics revealed that the decidualized microenvironment extends neutrophil survival and shapes discrete tissue-resident subsets, including an activated CD63^high^ phenotype. Imaging mass cytometry uncovered a precise spatial organization, demonstrating preferential cell-cell interactions between these activated neutrophils and invading EVTs. Furthermore, functional in vitro assays demonstrated that the neutrophil secretome, including the core granular protease neutrophil elastase, directly stimulates EVT invasion. Together, these findings identify decidua-associated neutrophils as crucial cellular coordinators of human placental development.

**One-sentence summary:** Neutrophil phenotype balance in early placentation shapes the EVT invaded decidua immune environment and drives trophoblast migration.

## INTRODUCTION

The establishment of the maternal-fetal interface during early human pregnancy requires a highly orchestrated cascade of tissue remodeling and immunological adaptation. Following blastocyst implantation into the endometrium, extravillous trophoblasts (EVTs) of fetal origin invade the maternal uterine stroma and spiral arteries to optimize uteroplacental blood flow (*1*). Driven by progesterone signaling and successful implantation, the endometrium undergoes decidualization to transform into structurally distinct zones in early pregnancy (*2*). The decidua basalis (DB) forms the anatomical site of extensive EVT invasion and direct placental attachment, whereas the decidua parietalis (DP) lines the remainder of the uterine cavity without placental contact, and the decidua capsularis (DC) transiently encapsulates the developing embryo (*3*). Dysregulation of these localized invasive processes is strongly linked to pregnancy complications, including recurrent miscarriage and preeclampsia (*4,5*).

While historical paradigms have focused heavily on the roles of adaptive T cells, decidual natural killer (dNK) cells, and tolerogenic macrophages in the decidua, a comprehensive, high-dimensional map of the decidual immunome remains incomplete. A major limitation of previous studies has been the reliance on bulk decidual tissue analysis. By failing to separate the DB from the DP, prior approaches have averaged the signals of heterogeneous cell populations, masking zone-specific activation states and cellular complexities (*6–8*). Crucially, in heavily invaded areas of the decidua, severe morphological alterations occur, characterized by compromised tissue integrity, eroded vessels, extravasal erythrocytes, and fibrin deposits previously described by us as remodeling lesions. In these areas, local lymphocyte and macrophage populations decline, yet neutrophils preferentially and almost exclusively accumulate (*9*). While recent work has begun to highlight the presence of specialized fibrinoid-associated neutrophils at the maternal-fetal interface (*10*), a comprehensive, multi-compartment mapping of decidual subset heterogeneity, local phenotypic polarization, and systemic-local neutrophil adaptation remains to be established.

Neutrophils are classically defined as short-lived, polymorphonuclear granulocytes that serve as first responders to acute inflammatory insults through degranulation, reactive oxygen species (ROS) production, and neutrophil extracellular trap (NET) formation (*11*). Emerging evidence, however, highlights substantial phenotypic and functional plasticity within the neutrophil lineage, pointing to specialized roles in non-infectious tissue homeostasis and pathology (*12–14*).

Accumulating evidence from animal experiments highlights the functional versatility of neutrophils during gestation. In murine models of early pregnancy, targeted neutrophil depletion led to severe compromises in vascular remodeling and placental architectures, as well as in increased rates of fetal resorption, establishing these cells as active drivers of pregnancy success rather than passive bystanders (*15,16*). Parallel to these localized tissue findings, systemic profiling of peripheral blood from pregnant individuals reveals a characteristic gestational neutrophilia (*17*). This systemic remodeling is characterized by the expansion of low-density neutrophils (LDNs) and polymorphonuclear myeloid-derived suppressor cells (PMN-MDSCs), which exhibit distinct immunosuppressive and regulatory functions necessary to maintain systemic maternal-fetal tolerance (*18–20*).

Despite these insights from animal models and peripheral circulation, a major knowledge gap remains regarding the spatial distribution, phenotypic heterogeneity, and functional specialization of neutrophils within the human maternal-fetal interface during the critical first trimester. Prior human decidual characterizations have heavily focused on lymphocytes and classical macrophages, leaving the local neutrophil compartment profoundly underrepresented. This gap is largely driven by technical limitations: neutrophils possess exceptionally low baseline transcript content, causing them to be consistently masked or excluded in standard single-cell RNA-sequencing (scRNA-seq) datasets (*21*). Furthermore, conventional isolation workflows involving density centrifugation or freeze-thaw cycles frequently deplete or artifactually activate tissue-resident granulocytes (*22*), while a historical lack of validated surface markers has hindered the robust classification of specific tissue-associated neutrophil polarization states (*23*). Crucially, by failing to separate the physically adjacent but functionally distinct DB and DP, previous human studies have averaged out localized signaling patterns, leaving it unclear whether tissue-infiltrating neutrophils represent a site-specific, microenvironment-driven activation state.

By integrating 28-marker flow cytometry, multiplexed imaging mass cytometry (IMC), and publicly available single-cell transcriptomic sub-clustering, we reveal the specialized architecture of decidua-associated neutrophil subsets. We demonstrate that the decidualized uterine microenvironment actively shapes distinct, hyper-activated neutrophil subsets that preferentially track to structural remodeling lesions and areas of heavy EVT invasion. Ultimately, through functional *in vitro* modeling, we show that these localized neutrophil populations directly orchestrate the maternal-fetal interface by actively stimulating EVT invasion, identifying decidua-associated neutrophils as essential cellular coordinators of human placental development.

## RESULTS

### High-dimensional single-cell profiling reveals a distinct myeloid shift and profound neutrophil enrichment in the decidua basalis

To characterize the local immune topography of the maternal-fetal interface during early gestation, we implemented a rigorous tissue-separation protocol focusing on localized remodeling lesions as previously described by us (*9*). Because macroscopically distinguishing the decidua basalis (DB) from adjacent decidual compartments like the decidua parietalis (DP) can be imprecise due to the lack of clear anatomical boundaries and extensive site remodeling, we applied an EVT-based functional definition to isolate decidual compartments. Specifically, tissues were separated based on the presence of remodeling lesions and classified as EVT-invaded decidua (iDec) versus non-invaded decidua (niDec), rather than relying solely on traditional DB and DP designations. Flow cytometry was used to validate tissue identity and purity, by EVT presence (CD45^-^ HLA-G^+^), which accounted for approximately 5% of total viable cells, whereas EVTs were completely undetectable in matched niDec samples (Fig. 1A). Additionally, the total immune cell infiltration (CD45^+^) was significantly higher in the iDec compared to the niDec, accounting for a mean of roughly 88% versus 75% of total live cells, respectively (Fig. 1B). Of note, these high baseline CD45^+^ percentages reflect an enzymatic isolation protocol optimized specifically for leukocyte recovery, which inherently skews the ratio of non-immune (CD45^-^) to immune (CD45^+^) cells toward an overrepresentation of leukocytes. Nevertheless, applying this validated tissue-digestion protocol ensures the comprehensive extraction and recovery of the complete decidual immune spectrum (*24*). To map the immunological differences between these adjacent microenvironments, we designed 28-marker flow cytometry panels, resolving 33 discrete immune (sub)populations and activation states (Table S1, fig. S1). Unsupervised visualization of the main immune cell lineages via sunburst plots revealed a striking divergence in the overall immune composition between the two adjacent decidual compartments (Fig. 1C). The iDec microenvironment exhibited a profound shift toward myeloid dominance. Although regulatory NK cells, M2 macrophages (CD68^+^CD206^+^), and CD14^+^HLA-DR^+^ monocytes remained present, the architectural landscape was overwhelmingly defined by a massive expansion of neutrophils. In contrast, the niDec microenvironment was heavily skewed toward the lymphoid lineage, dominated by natural killer (NK, CD3^-^CD56^+^) cells. Subpopulation analysis showed that these were primarily regulatory dNK cells (CD3^-^CD56^+^CD16^-^), alongside smaller fractions of CD16^+^ NK cells, with CD3^+^ T cells (comprising CD4^+^ and CD8^+^ fractions) making up the next largest lymphoid pool (Fig. 1C). To rigorously quantify these compositional shifts, we evaluated the relative abundance of the major leukocyte populations as a percentage of total CD45^+^ cells (Fig. 1D) and live cells/mg tissue (fig. S2). This cross-compartment comparison confirmed that NK cells represented the single largest leukocyte population in the niDec, contributing up to 70% of all leukocytes, but dropped significantly to low baseline percentages in the iDec.

**Fig. 1.**
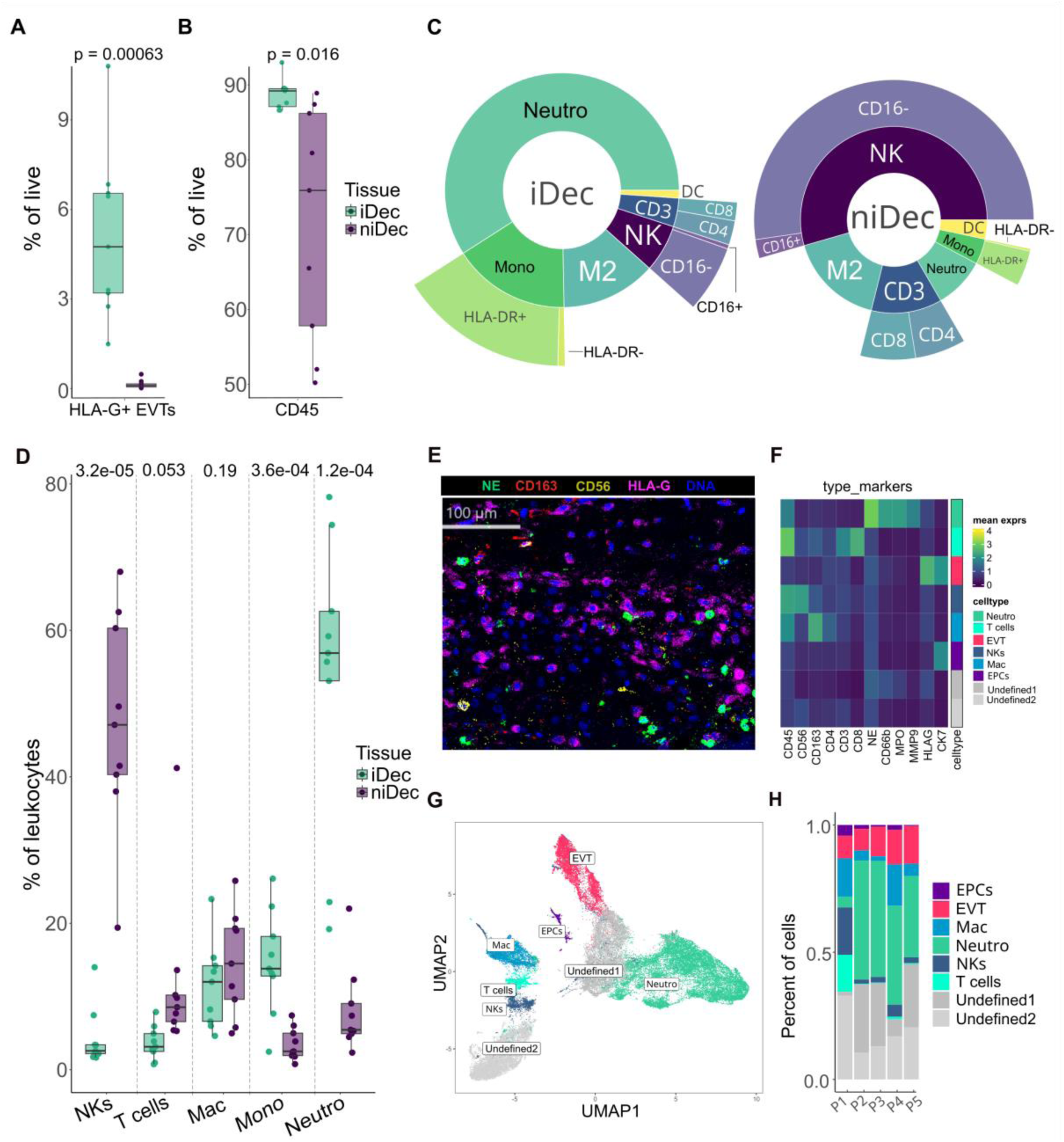
Neutrophils are the dominant immune cells in iDec. (**A**) % live CD45-HLA-G+ cells in iDec and niDec (N=9). (**B**) % live CD45+ cells in iDec and niDec (N=9). (**C**) Sunburst plot illustrating main immune populations and subpopulations within iDec and niDec. (**D**) Main immune populations in iDec and niDec represented as a % of live leukocytes (N=9). (**E**) Representative image depicting the expression of immune phenotyping markers in iDec tissue sample. Neutrophils (NE) are shown in green, macrophages (CD163) in red, NKs (CD56) in yellow, and EVTs (HLA-G) in magenta. Cell nuclei are depicted in blue. (**F**) Heatmap representing mean expression of type markers used for cluster annotations. (**G**) UMAP representation of immune cells present in iDec, obtained with IMC (N=5). (**H**) Abundance of different cell types in each iDec sample, presented as percentage of all cells. Paired Student’s t-tests were conducted to compare the two groups. Data are presented as mean ± SD, *P < 0.05.

Consistently, Solders *et al.* reported minimal overall NK cell enrichment in the iDec (*25*). Neutrophils inversely mirrored the NK distribution, accumulating massively in the iDec where they formed the predominant leukocyte population, averaging approximately 60% of total leukocytes, compared to an almost complete absence in the niDec. Comparable neutrophil numbers and high relative abundances within first-trimester DB tissue have been reported previously (*26*). Monocytes showed a significant enrichment in the iDec relative to the niDec, making up roughly 15% of the leukocyte pool in iDec. Macrophages and T Cells remained evenly balanced, exhibiting no statistically significant differences in relative frequencies between the iDec and niDec microenvironments.

To independently validate these findings and preserve native tissue architectures, we performed imaging mass cytometry (IMC) on formalin-fixed, paraffin-embedded (FFPE) first-trimester iDec tissue sections (Fig. 1E–H). Imaging captured a high density of iDec-associated neutrophils, identified by neutrophil elastase (NE) expression, distributed near other immune cells and EVTs (Fig. 1E). Cell annotation was performed based on the mean expression of cell type-specific markers (main immune cell differentiation markers). Neutrophils express several neutrophil-associated markers, such as NE, CD66b, MPO, and MMP-9. Other CD45^+^ immune cells were annotated based on the expression of CD56 (NKs), CD163 (macrophages), and CD3 (T cells). Both epithelial cells (EPCs) and EVTs express cytokeratin 7 (CK7) and were separated by HLA-G expression (*27*). Two distinct clusters did not express any marker (Fig. 1F). Uniform Manifold Approximation and Projection (UMAP) embedding of the IMC dataset confirmed the topological separation of these major lineages (Fig. 1G). Spatial quantification across multiple patient donor blocks confirmed that neutrophils consistently represent the dominant infiltrating immune population within the early iDec, confirming our flow cytometric findings (Fig. 1H).

### The decidualized tissue microenvironment modulates neutrophil activation and effector functions

Neutrophils represent the primary innate immune responders to chemoattractant gradients, rapidly transmigrating from peripheral circulation into target tissues (*28*). Within the iDec, neutrophils constitute the predominant infiltrating immune cell population, accounting for approximately 50% of total viable cells (Fig. 2A), amidst a cellular microenvironment rich in fibroblasts, decidual, stromal, epithelial, endothelial, and EVT cells (*29*). Upon activation, neutrophils undergo degranulation, releasing specialized granular proteins, including myeloperoxidase (MPO), matrix metalloproteinase-9 (MMP-9), and neutrophil elastase (NE), into the extracellular space (*30*). Indicative of this process *in situ*, we detected a significant positive correlation between the magnitude of decidua-associated neutrophils and the accumulation of MPO and MMP-9 within matched tissue explant supernatants (Fig. 2B). These findings align with our previous observations demonstrating elevated MPO expression in DB supernatants relative to DP counterparts (*31*).

**Fig. 2.**
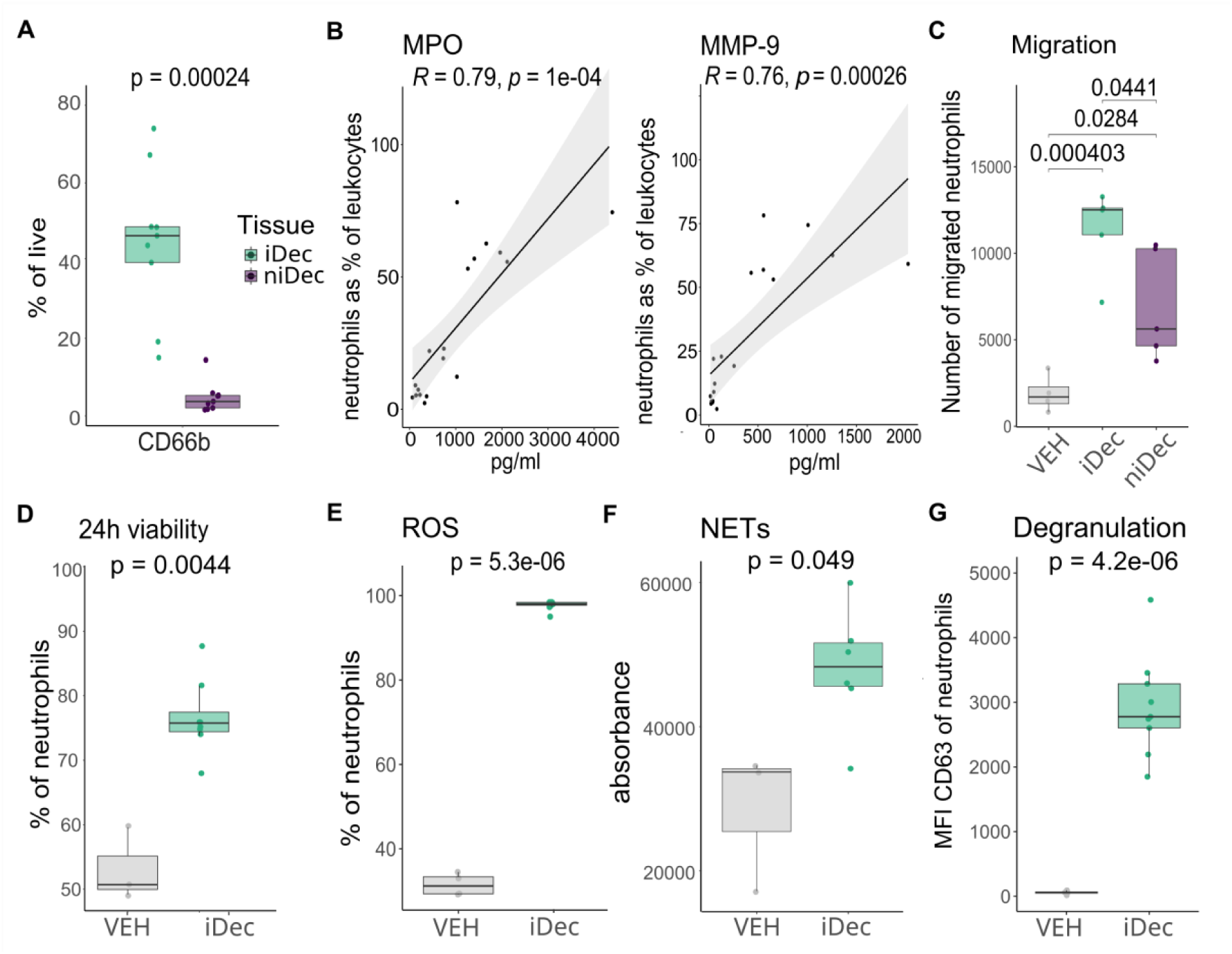
The tissue microenvironment modulates neutrophil activation and effector functions. (**A**) Neutrophil infiltration in iDec and niDec presented as % of all live cells (N=9). (**B**) Correlation of neutrophil infiltration presented as % of all leukocytes with amount of MPO (left) and MMP-9 (right) in tissue samples. (**C**) Healthy blood neutrophil migration stimulated with VEH, iDec, and niDec tissue explant supernatants (N=5). (**D**) Viability after 24h incubation of healthy neutrophils stimulated with VEH or iDec tissue explant supernatants (N=8). (**E**) ROS production by healthy neutrophils stimulated with VEH or iDec tissue explant supernatants (N=9). (**F**) NETs production of healthy neutrophils stimulated with VEH or iDec tissue explant supernatants (N=6). (**G**) Degranulation represented as CD63 MFI expression of healthy neutrophils stimulated with VEH or iDec tissue explant supernatants (N=8). Paired Student’s t-tests were conducted to compare two groups within the tissues of the same patient. Unpaired Student’s t-tests were conducted for comparing two groups. Pearson’s correlation analysis was performed. ANOVA with Tukey’s post-hoc test was performed to compare multiple groups. Data are presented as mean ± SD, *P < 0.05.

To elucidate the specific mechanisms by which the local uterine microenvironment shapes neutrophil behavior, we functionally evaluated peripheral blood neutrophils isolated from non-pregnant healthy donors following stimulation with tissue-derived supernatants, using vehicle-treated (VEH) neutrophils as a baseline control. Given that ILC3 cells residing in the DB are known to secrete the potent neutrophil chemoattractant IL-8 (*32*), we assessed directional migration using transwell experiments. Distinctly elevated neutrophil chemotaxis was observed toward tissue explant supernatants compared to VEH controls, with iDec supernatants eliciting a significantly more robust migratory response than niDec supernatants (Fig. 2C). Because niDec-induced migration was less prominent, subsequent functional analyses focused exclusively on the regulatory dynamics within the iDec microenvironment.

While the physiological lifespan of circulating neutrophils is restricted to several hours, inflammatory activation is known to prolong their survival over multiple days (*33*). We therefore investigated whether exposure to the iDec microenvironment could extend neutrophil viability. Whereas control blood neutrophils exhibited approximately 50% viability after 24 hours of *in vitro* culture, stimulation with iDec tissue supernatants significantly sustained viability, with nearly 80% of cells remaining viable at the 24-hour mark (Fig. 2D). Beyond enhanced survival, the iDec microenvironment profoundly augmented neutrophil effector functions. Supernatant-stimulated neutrophils exhibited robust reactive oxygen species (ROS) production, contrasting sharply with baseline conditions where only ∼20% of control neutrophils generated ROS (Fig. 2E). Furthermore, exposure to iDec supernatants significantly accelerated neutrophil extracellular trap formation (NETosis) relative to controls (Fig. 2F). Together with the observed prolongation of neutrophil viability, these findings suggest that iDec-derived soluble factors may promote a NET-prone yet sustained neutrophil state, potentially involving vital NETosis. However, this interpretation requires further validation, particularly through in vivo assessment of NET formation and neutrophil fate at the maternal-fetal interface. Finally, we quantified the surface expression of CD63, a sensitive marker of azurophilic granule exocytosis (*34*). Stimulated neutrophils displayed a striking increase in CD63 mean fluorescence intensity (MFI) compared to VEH controls (Fig. 2G). Collectively, these data demonstrate that the decidual tissue microenvironment serves as a critical determinant of neutrophil viability, degranulation, and oxidative effector programs.

### Phenotypic segregation and subset dynamics of tissue versus peripheral blood neutrophils

Given the high abundance of neutrophils infiltrating the iDec and the scarcity of data regarding their functional specialization, we characterized the surface phenotyping profiles of tissue and peripheral blood neutrophils. To rule out the possibility that the observed tissue-associated neutrophils resulted from maternal peripheral blood contamination, we performed principal component analysis (PCA) utilizing a panel of key immunophenotypic markers: CD62L, CD10, CD11b, CD63, and CXCR4. The PCA plot demonstrated distinct segregation between peripheral blood and decidua-associated neutrophils, suggesting distinct phenotypic profiles between these compartments (Fig. 3A). These findings support the notion that decidua-associated neutrophils represent a locally adapted phenotype. While neutrophils from the iDec and niDec also expressed divergent marker profiles, they exhibited closer phenotypic proximity to each other than to their peripheral counterparts (Fig. 3A).

**Fig. 3.**
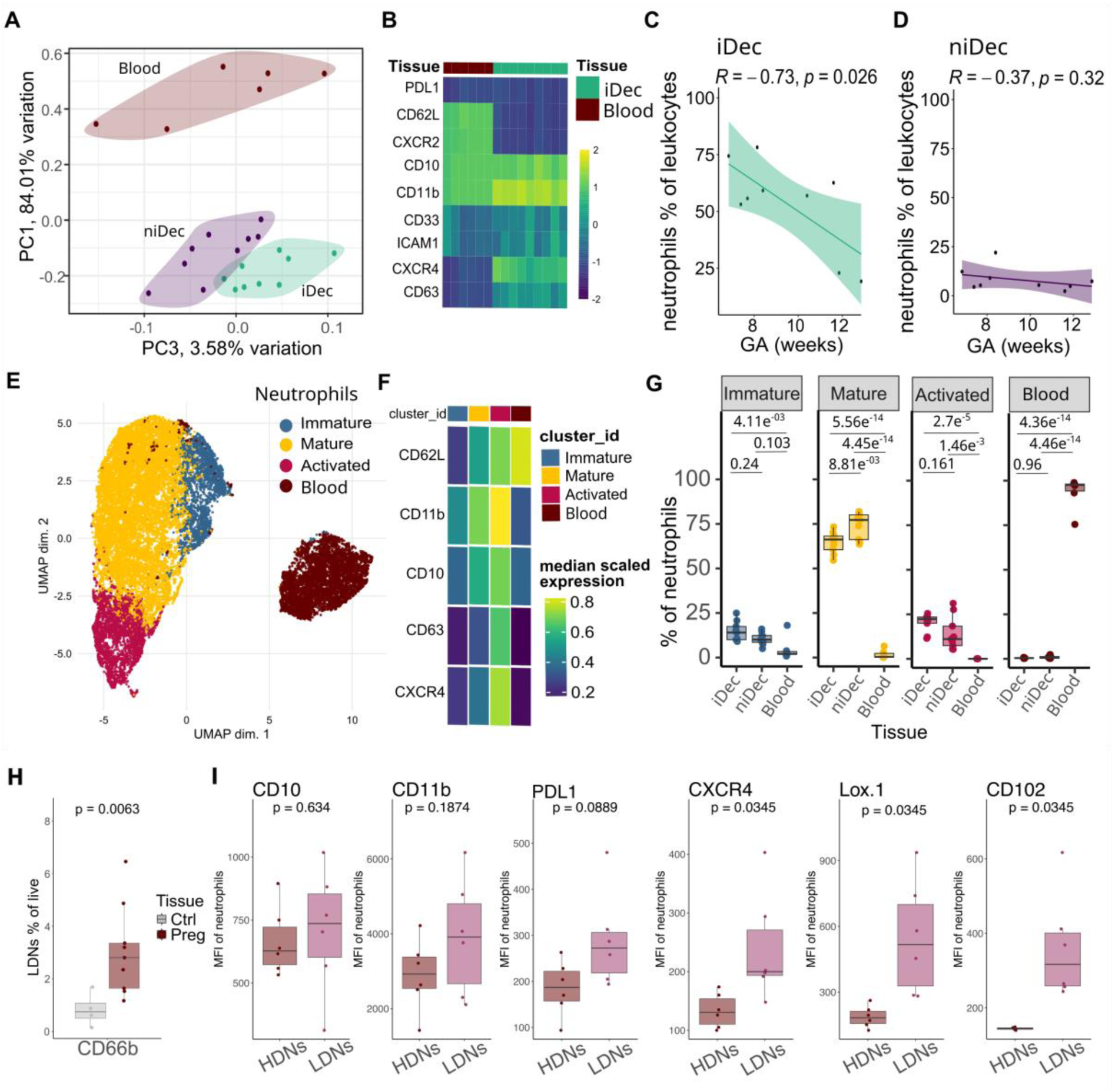
Phenotypic segregation of neutrophil subsets in tissues and blood. (**A**) PCA plot of neutrophils in blood samples from pregnant women, iDec, and niDec from elective abortion samples, compared based on phenotyping markers. (**B**) Heatmap of neutrophil phenotyping markers presented as % expression of neutrophils. Correlation of neutrophil infiltration in (**C**) iDec and (**D**) niDec with gestational week. (**E**) UMAP representing clustering of iDec, niDec, and blood neutrophils into four distinct clusters. (**F**) Heatmap of markers used for clustering, and their expression in each cluster. (**G**) Presence of each cluster within iDec, niDec, and blood neutrophils. (**H**) Presence of LDNs in PBMC fractions of non-pregnant and pregnant blood samples. Data is represented as % of live cells (N=9). (**I**) Comparison of neutrophil phenotyping markers on LDNs and HDNs within blood samples from pregnant women (N=6). Pearson correlation was performed to investigate relationship between variables. Unpaired Student’s t-test was conducted for comparing LDNs presence in non-pregnant and pregnant individual’s blood. Paired Student’s t-tests were conducted for comparing two groups from the same patient. ANOVA with Tukeys post-hoc test was performed for comparing multiple groups. Data are presented as mean ± SD, *P < 0.05.

We next analyzed the activation status of peripheral blood and iDec-associated neutrophils, quantifying scaled protein expression via heatmap analysis (Fig. 3B). Extravasation and infiltration into the decidual tissue were accompanied by a marked downregulation of CD62L and CXCR2 expression, a shedding pattern characteristic of tissue-transmigrated neutrophils (*35*). Both tissue-resident and peripheral populations maintained high expression of CD10, establishing their mature status. However, decidua-associated neutrophils exhibited significant upregulation of CD11b, CXCR4, and CD63, validating an advanced state of local activation (Fig. 3B). Expression patterns of established neutrophil markers varied across neutrophil populations in both iDec and niDec (fig. S4).

To determine whether neutrophil recruitment is temporally regulated during early gestation, we performed correlation analyses between iDec-associated neutrophils and gestational week (GW). Within iDec tissue specimens, we detected a significant negative correlation between the magnitude of neutrophil infiltration and GW, underscoring a precise temporal requirement for these cells during the critical implantation and early placental development phases (Fig. 3C). In contrast, neutrophil abundance within the niDec remained uniform across the evaluated first-trimester time points, showing no temporal correlation (Fig. 3D).

### The decidual microenvironment shapes discrete neutrophil subsets and favors an activated phenotype

Distinct neutrophil subpopulations have previously been delineated in circulation and peripheral tissues based on discrete surface protein expression patterns (*23,36,37*). To interrogate the phenotypic heterogeneity within the iDec, niDec, and peripheral blood, we performed unsupervised clustering analysis utilizing the marker panel (CD62L, CD10, CD11b, CD63, and CXCR4). UMAP analysis identified three phenotypically distinct decidua-associated neutrophil clusters, which we classified as: immature (CD10^-^), mature (CD10^+^) and activated (CD63^high^). Consistent with our PCA analysis, peripheral blood neutrophils formed a separate, distinct cluster from the decidua-associated neutrophils (Fig. 3E). Median-scaled expression analysis identified CD62L expression as the primary driver segregating the peripheral blood population. Within the decidual tissues, the activated subset exhibited globally elevated MFIs across all analyzed markers, indicative of a hyper-activated phenotype (Fig. 3F). To assess how the specific uterine microenvironment influences these subsets, we quantified cluster frequencies within each tissue compartment, normalizing cell inputs to account for variations in absolute neutrophil counts. The activated subset comprised approximately 25% of the total neutrophil pool in the iDec, compared to roughly 15% in the niDec (Fig. 3G). Notably, activated neutrophils were absent from peripheral blood, further ruling out systemic contamination of the tissue samples. While the iDec also contained a higher frequency of immature neutrophils than the niDec, this trend did not reach statistical significance. Conversely, mature neutrophils were more enriched in the niDec than in the iDec, demonstrating that the localized immune microenvironment of the maternal-fetal interface actively drives distinct phenotypic skewing (Fig. 3G).

### Phenotypic characterization of pregnancy-associated low-density neutrophils

Finally, because healthy pregnancy is characterized by systemic neutrophilia (*17*), we evaluated whether circulating neutrophil subsets are simultaneously modulated. Low-density neutrophils (LDNs) have been documented to expand during gestation (*20*); thus, we compared the frequency of peripheral LDNs between first trimester pregnant individuals and non-pregnant controls. Our analysis revealed a significant expansion of circulating LDNs during pregnancy, averaging 3% of total viable cells, compared to less than 1% in non-pregnant controls (Fig. 3H), indicating that gestational status drives systemic innate immune remodeling alongside localized decidual polarization.

To further define the immunological properties of the expanded circulating LDN population during pregnancy, we compared their surface marker profile against autologous high-density neutrophils (HDNs). No significant differences were observed between LDNs and HDNs in the expression of the maturation marker CD10, the activation/checkpoint marker PD-L1, or the activation marker CD11b. However, LDNs displayed a distinct phenotypic shift characterized by the significant upregulation of CXCR4 (a chemokine receptor implicated in tissue homing and retention), LOX-1 (a lectin-like oxidized LDL receptor-1), and CD102 (ICAM-2, an intercellular adhesion molecule) (Fig. 3I). Elevated CXCR4 expression has previously been identified on LDNs during severe sepsis, where it promotes tissue retention and prolongs cellular survival (*38*). LOX-1 has emerged as a surface marker distinguishing immunosuppressive polymorphonuclear myeloid-derived suppressor cells (PMN-MDSCs) within the LDN fraction (*39*). Taken together, these phenotypic signatures indicate that pregnancy-associated circulating LDNs possess an activated, tissue-tropic profile. This phenotype may facilitate their targeted extravasation into gestational tissues, where they could actively contribute to localized maternal-fetal immunomodulation.

### Spatial and single cell profiling identifies conserved neutrophil sub-populations within the iDec

Neutrophils are increasingly recognized as pivotal orchestrators of both adaptive and innate immune dynamics across a spectrum of physiological and pathological contexts (*40,41*). Given that immune tolerance and tissue remodeling during early gestation rely heavily on coordinated cross-talk between maternal leukocytes and fetal EVTs (*42–44*), we leveraged IMC to interrogate the microenvironmental architecture and spatial distribution of HLA-G^+^ EVTs and iDec-associated neutrophils. Imaging of neutrophil proteins, including CD66b, MPO, NE, and MMP-9, revealed a striking spatial proximity between neutrophils and invading EVTs, prominently highlighted within highly magnified tissue areas (Fig. 4A). By extracting these discriminatory markers, we successfully segregated neutrophils into a distinct cluster, as illustrated in Fig. 1G, demonstrating conserved expression patterns within this cluster on UMAP plots (Fig. 4B).

**Fig. 4.**
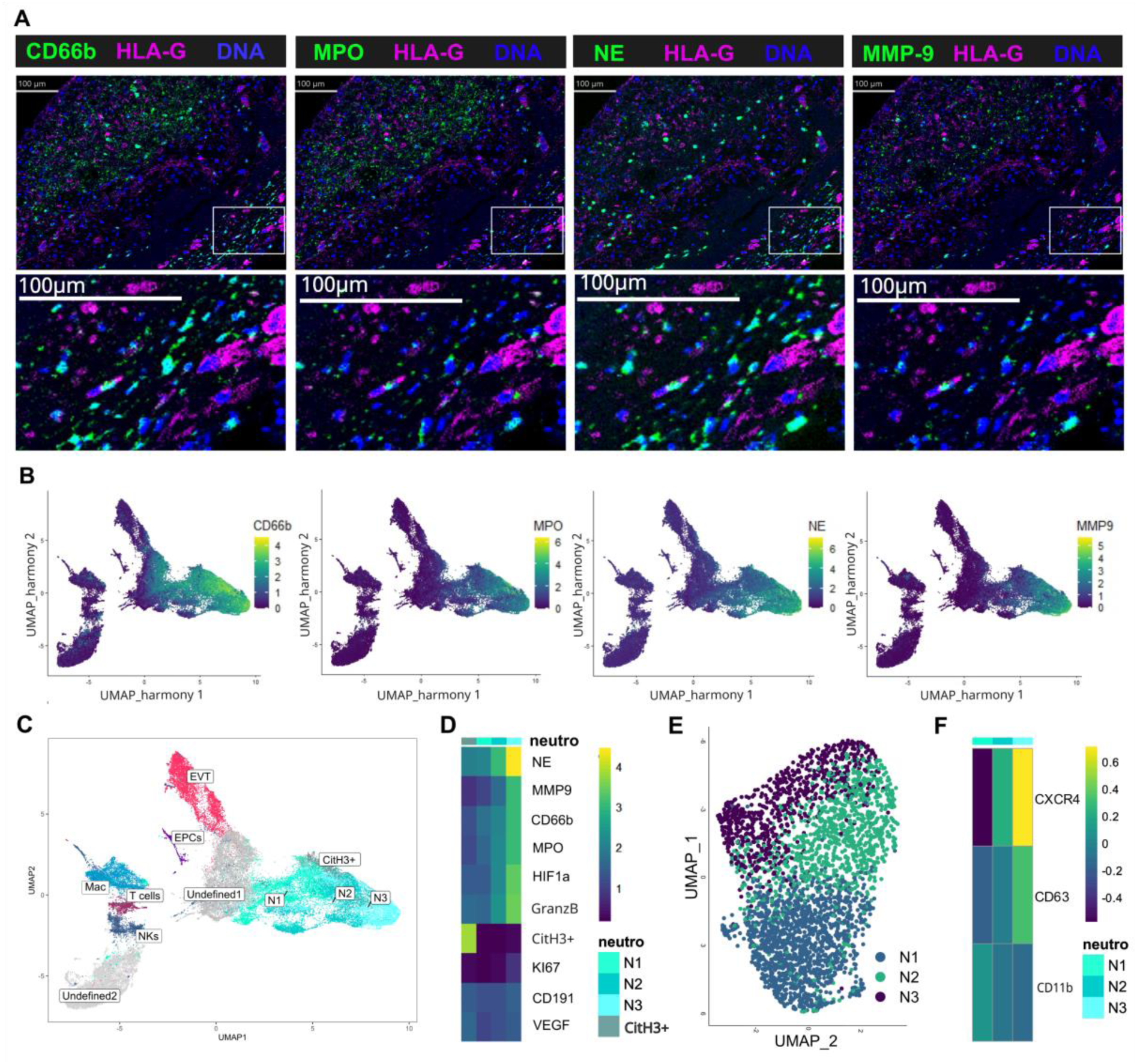
Different neutrophil subsets within the tissue. (A) Representative IMC images of neutrophil associated markers (green) and EVTs as HLA-G^+^ cells (magenta). Nucleus is depicted in blue. Lower panels show magnified view of selected spot. (**B**) UMAP representations of expression of neutrophil associated markers. (**C**) UMAP showing neutrophil distinct clusters in combination with other cell types. (**D**) Heatmap of markers used for neutrophil subclustering, and their expression in each cluster. (**E**) UMAP showing neutrophil clusters in scRNAseq datasets. (**F**) Heatmap showing gene clustering of publicly available scRNAseq DB neutrophils (*45*).

To determine whether the cellular heterogeneity observed in our flow cytometry data was recapitulated by IMC, we further subclustered neutrophils based on the main neutrophil granular proteins in combination with several activation markers. Within the tissue, four different clusters of neutrophils were found, denoted as N1, N2, and N3, and a smaller subset expressing CitH3 was indicative of NETotic neutrophils (Fig. 4C). These clusters exhibit differential expression of NE, MMP-9, CD66b, MPO, HIF1α, and Granzyme B, with cluster N3 displaying the highest expression of these proteins, suggesting potential activation of these cells. Despite the technical use of distinct marker configurations for flow cytometry and IMC analysis (Table S2), we observed parallel activation profiles and phenotypic trends across both independent protein-detection platforms (Fig. 4D).

We next investigated whether these phenotypically defined intra-tissue sub-populations could be validated at the transcriptional level. We interrogated a publicly available first-trimester DB single-cell RNA sequencing (scRNA-seq) dataset (*45*) to profile iDec-associated granulocytes. Because classic neutrophil-associated transcripts are often captured at low depths in multiplexed tissue architectures, standard sub-clustering of the scRNA-seq dataset was focused exclusively on the consistently detected loci CXCR4, CD63, and ITGAM (encoding CD11b), which mirrored the primary surface markers utilized in our initial flow cytometry experiments (Fig. 4E). Unsupervised transcriptional sub-clustering of this scRNA-seq granulocyte fraction successfully resolved three distinct neutrophil clusters, each corresponding to a unique molecular activation state (Fig. 4F). Taken together, the identification of three primary neutrophil sub-populations across three independent experimental methodologies, suggests the presence of a structured, non-random landscape of specialized neutrophil phenotypes at the maternal-fetal interface during early pregnancy.

### Spatial proximity and cell-cell interaction analysis reveal preferential neutrophil-EVT crosstalk

To decode the architectural connectivity and potential crosstalk between specific decidual neutrophil sub-populations and neighboring maternal or fetal cell types, we conducted spatial interaction analyses on our IMC dataset. Spatial neighborhood modeling demonstrated that neutrophils are non-randomly organized relative to other cell types within the iDec. Notably, the phenotypically mature and activated neutrophil subsets, designated as N2 and N3, were predominantly localized in immediate proximity to infiltrating HLA-G^+^ EVTs, as highlighted within the representative spatial region (Fig. 5A). To quantify these spatial relationships, we performed patch interaction analysis to determine the significance of cellular proximity. This computational modeling revealed that EVTs preferentially interact with the N2 and N3 neutrophil subsets over other immune or structural cells, with the highest density of positive spatial interaction scores tracking specifically to the hyper-activated N3 subset (Fig. 5B).

**Fig. 5.**
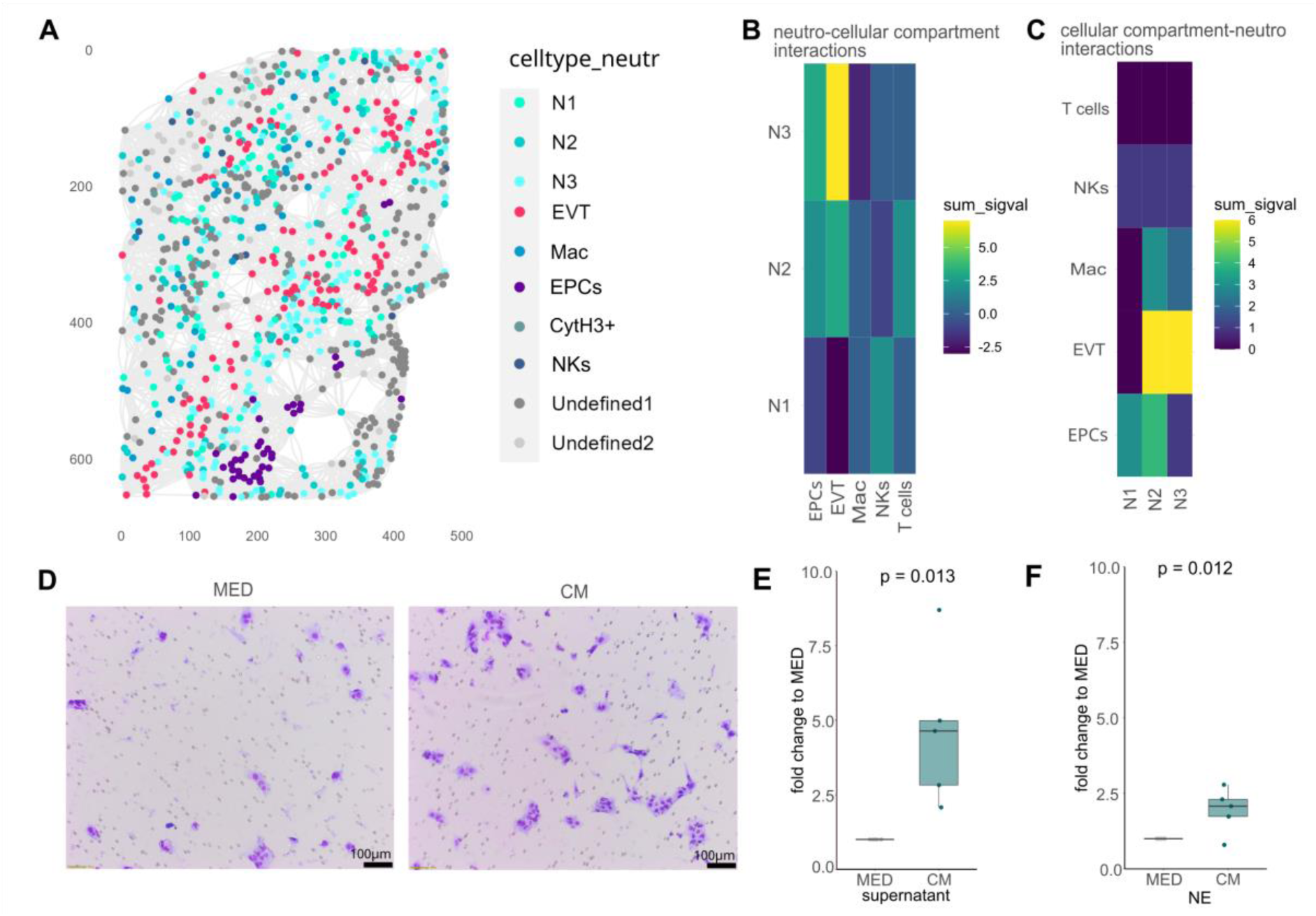
Neutrophils interact with EVTs and stimulate their invasion. (**A**) Representative k-nearest neighbors (KNN) graph showing localization and interactions of neutrophil subcluster with other cell types. (**B**) Heatmaps showing patch interaction analysis between cell types and neutrophil subsets. (**C**) Heatmaps showing patch interaction analysis between neutrophil subsets and other cell types. (**D**) Representative images of JEG-3 invasion upon stimulation with control media (MED, left) and neutrophil supernatants (CM, right). (**E**) Fold change of invaded JEG-3 cells stimulated with blood neutrophil supernatants (CM) compared to media (MED) (N=5). (**F**) Fold change of invaded JEG-3 cells stimulated with NE compared to MED (N=5). Student’s t-tests were conducted for comparing two groups. Data are presented as mean ± SD, *P < 0.05.

Reciprocally, looking from the neutrophil perspective, the N2 and N3 subsets exhibited highly elevated and comparable interaction signals centered on EVTs relative to their crosstalk with local maternal T cells, NKs, macrophages, or EPCs (Fig. 5C). Collectively, these topological findings demonstrate an intricate, spatially restricted interplay between advanced effector neutrophil subsets and invading EVTs at the placental interface.

### Decidual neutrophils actively promote extravillous trophoblast invasion *in vitro*

Because our spatial analyses revealed a tight topographical association between activated neutrophils and invading EVTs, we sought to determine the functional consequences of this cellular crosstalk. Given the vital role of adequate EVT invasion in driving spiral artery remodeling, we investigated whether secretome components derived from activated neutrophils directly modulate EVT invasion. Using the established human EVT-surrogate JEG-3 choriocarcinoma cell line as an invasion model, as previously described (*31*), we performed transwell invasion assays. JEG-3 cells were stimulated with conditioned supernatants harvested from healthy peripheral blood donor neutrophils or NE as condition media (CM) or treated with empty media (MED). We observed a significant increase in the number of invaded JEG-3 cells following exposure to neutrophil-derived supernatants and NE compared to vehicle-treated wells (Fig. 5D-F).

Taken together, these functional data complement our spatial findings, demonstrating that iDec-associated neutrophils and their secretory factors serve as active modulators within the decidual microenvironment to facilitate adequate EVT invasion and support normal morphogenesis of the maternal-fetal interface.

## DISCUSSION

The early stages of human placentation demand a highly orchestrated balance between structural tissue remodeling, robust vascular modification, and localized immunotolerance (*1,2*). While the current framework has attributed these maternal-fetal adaptations predominantly to dNKs and macrophages (*6,25,29*), our study provides a comprehensive, high-dimensional map that identifies neutrophils as immunological architects of the first-trimester iDec. By separating the physically adjacent but functionally distinct iDec and niDec, we uncover a striking myeloid shift unique to the site of active fetal invasion, demonstrating that the decidualized microenvironment actively reprograms neutrophil subsets to drive EVT motility.

Our flow cytometry and IMC analyses reveal a profound architectural divergence across the decidual landscape: whereas the non-invaded Dec remains a lymphoid niche dominated by regulatory dNK cells, the heavily invaded Dec is overwhelmingly dominated by neutrophils, comprising over half of the local leukocyte pool. This high granulocytic infiltration directly challenges traditional views of the first-trimester immunome, which have frequently underrepresented neutrophils due to technical exclusions in standard scRNA-seq workflows or artifactual depletion during density-gradient processing or sample freeze-thaw protocols (*21,22*). While recent spatial analyses and related work have begun to highlight the presence of neutrophils at the maternal-fetal interface (*9,46,47*),our study reveals that this infiltration is neither random nor uniform, but is defined by compartmental restriction and precise cellular topography.

This localized accumulation is driven by active microenvironmental recruitment and polarization. The human placenta establishes zone-specific immunity by fundamentally remodeling the local DB secretome; specifically, DB-associated macrophages undergo an EVT-driven polarization that prompts the high-level secretion of neutrophil-priming and chemoattractant factors like CXCL1 and CXCL5 (*26*). Further, ILC3-derived IL-8 in the DB (*32*), exerted a potent chemotactic gradient on circulating neutrophils. This highly specialized tissue secretome aligns with our current data demonstrating enhanced directional migration, prolonged viability, ROS production and rapid activation of healthy neutrophils when exposed to iDec explant supernatants.

Furthermore, this local signaling network is structurally concentrated within the heavily invaded areas of the DB characterized by compromised tissue integrity, eroded vessels, and fibrin deposits, representing a specialized microenvironment previously termed ’remodeling lesions’ (*9*). Concurrently, spatial neighborhood analyses demonstrate that N2 and hyper-activated N3 neutrophil subsets preferentially localize and interact directly with invading fetal HLA-G^+^EVTs within these remodeling lesions. Building upon our previous findings that MPO is significantly elevated in DB supernatants and actively internalizes into JEG-3 cells to stimulate invasion (*31*), our new functional assays reveal that both neutrophil-conditioned media and the granular proteases, NE directly drive EVT invasion. Together with recent evidence highlighting MMP-9 production by fibrinoid-associated neutrophils (*10*), these data establish that decidual neutrophils deploy a coordinated proteolytic secretome (MPO, NE, and MMP-9) that actively facilitates trophoblast motility and normal morphogenesis of the maternal-fetal interface.

Our data indicate that these decidua-associated neutrophils do not reflect passive peripheral blood contamination, but instead represent a highly plastic, site-specific population undergoing advanced localized adaptation. PCA and phenotypic profiling demonstrated that extravasation into the iDec prompts a coordinated surface remodeling program. This is marked by the shedding of CD62L and CXCR2, alongside the robust upregulation of CD11b, the homing marker CXCR4, and the degranulation marker CD63. This activation state is driven by localized tissue signaling.

Prior studies have noted the abundance of neutrophils within the decidua (*26,36,47*); however, the precise phenotypes and roles of these cells at the human placental interface remained unknown. Functional importance is given to this population by animal models, where early-pregnancy neutrophil depletion directly causes implantation failure and aberrant placentation (*15,48*). Rather than functioning as a homogeneous population within the tissue, unsupervised spatial and transcriptional clustering resolved three conserved decidua-associated neutrophil subsets: immature (CD10^−^), mature (CD10^+^), and activated (CD63^high^). This activated subset was heavily enriched within the iDec compared to the niDec and completely absent from peripheral circulation. Strikingly, we observed a strict temporal regulation of this myeloid landscape, characterized by a significant negative correlation between iDec-associated neutrophil abundance and gestational age. This temporal restriction aligns with the peak window of trophoblast invasion and uterine remodeling, suggesting a highly synchronized physiological window of action.

Beyond localized neutrophil polarization within the decidua, we observed a parallel systemic adaptation in the maternal peripheral blood, marked by a significant expansion of circulating LDNs in pregnant individuals compared to non-pregnant controls. Phenotypic dissection reveals that these pregnancy-associated LDNs exhibit a unique profile defined by elevated expression of the retention marker CXCR4, the adhesion molecule CD102, and the immunosuppressive marker LOX-1. This suggests that specialized, immunosuppressive LDN subsets expand systematically to maintain maternal tolerance, while iDec-associated neutrophils undergo local activation under the influence of decidualizing cues, ultimately accelerating EVT-mediated vascular remodeling.

In conclusion, our study maps the spatial architecture, phenotypic heterogeneity, and functional specialization of first-trimester human decidual neutrophils, positioning them as essential cellular coordinators of early pregnancy success. By moving past historical bulk tissue limitations and profiling distinct sub-populations across independent experimental modalities, we define a structured, non-random landscape of myeloid-trophoblast crosstalk. Disruptions in this fine-tuned myeloid balance and spatial positioning may underpin the defective deep placentation seen in severe pregnancy complications such as preeclampsia and recurrent miscarriage. Unlocking the molecular targets within this newly defined decidua-associated neutrophil axis opens promising therapeutic horizons for immunomodulatory interventions in gestational disorders and reproductive success.

## MATERIALS AND METHODS

### Study design and approval

This study complied with the Declaration of Helsinki and was approved by the Ethics Committee of the Medical University of Graz (EK-number: 31-019 ex 18/19) and the Medical University of Vienna (EK-number: 084/2009). All participants provided written informed consent prior to enrollment in the Department of Obstetrics and Gynecology at the Medical University of Vienna (Austria) and the Department of Obstetrics and Gynecology at the Medical University of Graz (Austria). All elective terminations were performed by surgical vacuum aspiration and tissue was separated under a microscope, as previously described (*26*). Subsequently, iDec and niDec was defined by the presence (iDec) and absence (niDec) of HLA-G^+^ EVTs by flow cytometry. Blood samples were collected from pregnant women undergoing elective abortion (Vienna) or from pregnant women in their first trimester (Graz). The gestational age of the tissue and blood samples ranged from 6.9 to 13.7 weeks.

### Tissue processing

Tissues were collected, immediately processed, and stained for flow cytometry, as previously described (*24*). In brief, the iDec and niDec were microscopically separated, placed in RPMI1640 medium, and kept on ice. Subsequently, the tissue was washed in 40 ml of PBS without Ca^2+^ and Mg^2+^ (PBS −/−) for 2 min while continuous shaking. Any residual blood was removed, and the tissue was washed again with PBS. Small pieces were snap-frozen for further experiments and used for explant cultures. The remaining tissue was minced and further digested in RPMI-1640 supplemented with 40 U ml^−1^ DNAse I (Worthington) and 300 U ml^−1^ collagenase I (Worthington) at 37°C for 30 min. The tissue digest was homogenized by passing it through a 19-G needle and a 100-µm cell strainer. Digestion was stopped by adding 10% (v/v) fetal bovine serum (FBS). Erythrocytes were lysed using 1× RBC lysis buffer (BioLegend), and lysis was inactivated by adding four volumes of PBS−/−. Finally, cells were passed through a 50-µm cell strainer, and cell counts were determined using an EVE automated cell counter (NanoEntek).

### Blood processing

Donor blood was centrifuged at 100 g with a low brake for 20 min, and the plasma was carefully removed. Subsequently, 3 ml of 6% dextran was added and filled up to 25 ml with 0.9% sodium chloride solution, followed by incubation for 30 min at room temperature (RT). The upper phase of dextran sedimentation was placed on PBMC Spin Medium (PluriSpin) and centrifuged at 400 g with a low break for 20 min. PBMC and PMNL fractions were washed with HBSS. Lysis was performed using 10 ml of 0.2% saline for 30 s, followed by the addition of 10 ml of 1.6% saline. Cells were then washed with HBSS. Cell counts and viability were determined using an EVE automated cell counter.

### Immune phenotyping

Viability staining was performed immediately after generating the single-cell suspension, with staining performed in three separate panels. Cells were stained with a fixable viability dye (FVD) eFluorTM 780 (eBioscience) before fixation.

*Lymphocyte surface panel*: The cells were fixed for 10 min in IC fixation buffer (Invitrogen), washed in staining buffer (SB, PBS without Ca^2+^ and Mg^2+^ supplemented with 2% FBS), and stained on the following day. First, preincubation with an Fc receptor-blocking solution (BioLegend) was performed to reduce nonspecific binding. Subsequently, the cells were stained with fluorochrome-conjugated anti-human antibodies against HLA-G, CD45, CD3, CD4, CD8, CCR7, CD45RA, CD38, CD69, PD-1, CD19, CD20, CD56, and CD16 (for antibody and clone details, see Supplementary Table S1) for 30 min at 4°C and washed twice in SB.

*Lymphocyte intranuclear panel*: The cells were fixed for 10 min in 1× TF fixation buffer (BD), washed in SB, and stained on the following day. Before staining, the cells were fixed and permeabilized for 40 min using a BD transcription factor fixation/permeabilization kit to allow the antibodies to bind to the nuclear proteins. Subsequently, cells were preincubated with Fc receptor blocking solution and stained with fluorochrome-conjugated anti-human antibodies against CD45, CD3, CD4, and FoxP3 (for antibody and clone details, see Supplementary Table S1) for 30 min at 4°C. Cells were washed twice with TF washing buffer (BD Biosciences) and once with SB.

*Myeloid intracellular panel:* Cells were fixed for 10 min using the BD Cytofix kit, washed with BD Cytofix wash solution, and stained the following day. Before staining, the cells were again fixed and permeabilized for 20 min using the BD Cytofix/Cytoperm kit. Subsequently, cells were pre-incubated with Fc receptor blocking solution and stained with fluorochrome-conjugated anti-human antibodies against CD45, CD68, CD80, CD206, CD66b, Siglec 8, CD14, HLA-DR, CD16, CD117, CD11c, CD123, and PD-L1 (for antibody and clone details, see Supplementary Table S1) for 30 min at 4°C. Cells were washed twice with Cytofix wash (BD) and once with SB.

*Analysis:* All samples were analyzed using a BD LSRFortessa flow cytometer with FACSDiva software (BD Biosciences). Compensation was performed using single stains, and the background fluorescence cutoff values were based on the fluorescence minus one (FMO) strategy. Gating for each sample was based on an SSC height versus SSC area plot to eliminate aggregates. FVD staining was used to identify and eliminate dead cells. Analysis and compensation were performed using FlowJo software (TreeStar). For the sunburst plots, the percentages of cell types were rounded, and the R plotly package was used (*49*).

### Imaging mass cytometry (IMC)

Imaging mass cytometry was performed as previously described (*50*). Five iDec sections were stained, and multiple regions of interest (ROIs) were analyzed to cover the whole tissue section per sample. FFPE sections were deparaffinized by heating at 60°C and immersed in xylene twice for 10 min each. Decreasing alcohol concentrations (100%, 96%, 80%, 70%, and ddH_2_O for 5 min each) were used to rehydrate the samples. PBS-/-was used to wash the slides. Antigen retrieval (AR) involved incubating the samples in AR Reagent pH 9 for 30 min at 96°C (Agilent). Subsequently, when the slides were cooled, they were washed with PBS−/− for 10 min. Blocking was performed at RT for 45 min in 3% BSA (Sigma-Aldrich) in PBS−/− solution. The antibodies used for IMC are listed in Table S2. The first antibody cocktail was applied to the sections for 5 h at room temperature, followed by the second antibody cocktail at 4 °C overnight. After washing in 0.2% Triton-X 100 and PBS−/− for 8 min, tissue was stained with Intercalator-Ir diluted 1:1500 in PBS -/- for 5 min. The tissues were then washed with ddH2O for 5 min and air-dried. ROIs were measured using a Hyperion Imaging System (Standard BioTools) connected to a Helios mass cytometer (Standard BioTools). Samples were measured in two batches, and calibration was performed each time. The IMC data were analyzed using R software (v4.4.2). Initially, Docker with a Steinbock pipeline (v0.16.0) and Mesmer were deployed for efficient cell segmentation, as described previously (*51*). Subsequent analysis procedures followed the established Bodenmiller pipeline, ensuring a comprehensive and reliable examination of the IMC data (*52*) using the CATALYST (v1.30.2), imcRtools (v1.12.0), SpatialExperiment (v1.16.0), cytomapper (v1.18.0), bluster (v1.16.0), and SingleCellExperiment (v1.28.1) packages. Batch correction was performed using Harmony. A total of 56.583 cells was segmented across 5 iDec samples (7.714-14.986 cells per sample). Cell clustering was performed using ConsensusClusterPlus (v1.70.0) with FlowSOM-based clustering based on expression of the following markers: CD45, CD56, CD163, CD4, CD3, CD8, NE, CD66b, MPO, MMP9, HLA-G, and CK7. Spatial neighborhood analysis was performed using a k-nearest neighbor (kNN) approach implemented in imcRtools. The MCD Viewer software (v1.1.0) was used to visualize the antibody staining of the selected ROIs.

### Tissue supernatant generation and functional assays

Four small tissue fragments from different tissue sites (approximately 1mm^3^ each) per donor were incubated in 1 ml F12-DMEM without phenol red and supplemented with 1% penicillin/streptomycin for 24 h at 37°C in an atmosphere of 5% CO_2_. The supernatants were collected and snap-frozen for subsequent experiments.

*Viability:* A total of 200.000 PMNLs from healthy blood donors per sample were washed and resuspended in a 1:4 dilution of tissue supernatants in RPMI. Cells were incubated at 37°C under 5% CO_2_ for 24 h. Subsequently, cells were collected, washed with PBS-/- and stained with propidium iodide (PI) viability dye. Cell viability was immediately measured using a BD Canto flow cytometer with FACSDiva software (BD Biosciences).

*ROS:* PMNLs were incubated for 5 min in 5µM Dihydrorhodamine 123 and 5 µg/ml Cytochalasin B in PBS+/+ supplemented with 0.1% BSA, 10 mM HEPES, and 10 mM glucose. 50.000 cells per sample were incubated in a 1:4 dilution of the supernatants for 20 min at 37°C. Subsequently, cells were fixed (CellFIX diluted 10x in ddH_2_O, further diluted 4x in FACSFlow), and ROS levels were immediately measured using a BD Canto flow cytometer with FACSDiva software (BD).

*NETs:* PMNLS (2.67*10^6 cells/ml) were mixed with 1 drop/ml of SYTOX Green (BD, SYTOX™ Green Ready Flow™ Reagent). 200.000 PMNLs per well were mixed in a 1:4 dilution of tissue supernatants in RPMI or vehicle (PMA stimulation served as positive control) for 3 h at 37°C, 5% CO2 in a 96-well black plate (Thermo Fisher, Nunc™ F96 MicroWell™ Black and White Polystyrene Plate), each into two technical replicates. Absorbance was measured using a microplate reader (ClarioStar) at 488/520 nm. The mean value of each condition was subtracted from that of the unstained sample to eliminate background absorbance.

*Migration:* Neutrophil migration toward iDec and niDec supernatants was measured using 96-well transwell chambers with 5 µm polycarbonate membranes (Corning). Filters were pre-soaked in plus buffer (PBS with Ca2+/Mg2+ supplemented with 0.1% BSA, 10 mM HEPES, and 10 mM glucose) for 1 h at 37°C. PMNLs were isolated from healthy donors as described above. A total of 100 µl of supernatants diluted 1:2 with supernatant media were applied to the lower compartment, and 100 µl of PMNL suspension (4 million cells/ml) were added to the upper compartment. Cells were allowed to migrate for 1 h at 37°C and then fixed with 150 µL of fixative solution (CellFIX (BD) diluted 1:10 in distilled water with the addition of 1:4 FACSFlow (BD)). The number of migrated neutrophils was measured using BD FACSCanto II for 30 s. Neutrophils were separated from eosinophils based on their different autofluorescence in the V450 channel.

### ELISA

The ProcartaPlex 10-Plex ELISA kit (Thermo Fisher) was used for the quantification of MMP-9. A total of nine paired iDec and niDec tissue explant supernatants were used in this study. The supernatants were diluted 1:2, and the assay was conducted following the manufacturer’s protocol.

### MPO ELISA

Tissue was collected from elective first-trimester abortions and separated into iDec and niDec, as previously described (*26*). Tissue pieces were incubated at 37°C for 24 h in 0.5 ml F12/DMEM without phenol red media but supplemented with 1% P/S, and supernatant was collected. ELISA was performed and analyzed according to the manufacturers’ protocol (Human Myeloperoxidase DuoSet ELISA, R&D Systems, #DY3174). Data were initially published in (*31*) and re-analyzed for this study.

### Low density neutrophils

PBMC (containing LDNs) and PMNL (containing HDNs) fractions were stained using a flow cytometry surface-staining protocol. Neutrophils were first gated as SSC-A and FSC-A high cells. Furthermore, neutrophils (CD66b+, Siglec8-) were separated from eosinophils (CD66b-, Siglec8+ cells). MFIs of protein expression on neutrophils were analyzed to quantify the expression levels of specific surface proteins: CD10, CD11b, PD-L1, CXCR4, Lox-1, CD102. For antibody and clone details, see Table S1.

### Single-cell RNA sequencing data analysis

We reanalyzed a publicly available scRNA-seq dataset of first-trimester decidua (*45*), deposited at the National Omics Data Encyclopedia (www.biosino.org) under accession number OEP002901. The dataset contains decidual tissue from women with a normal pregnancy undergoing elective termination (n=5) and from patients with recurrent miscarriage (RM; n=10), at gestational age 7–9 weeks. As described by Bao et al. (*45*), single-cell libraries were prepared with the Chromium Single Cell Gel Bead Kit V3 (10x Genomics), sequenced on an Illumina NovaSeq 6000 (paired-end, 150 nt), and aligned and quantified against the GRCh38 human reference genome with CellRanger (V3). Downstream analyses were performed in R (v4.4.2) with the Seurat package (v5). Low-quality cells, defined as cells with fewer than 500 detected genes or more than 20% mitochondrial counts, were removed. Doublets were identified and removed with DoubletFinder (V2.0.3) (McGinnis et al., 2019). DoubletFinder was run on each sample separately. For each sample, the optimal neighborhood size (pK) was the one that maximized the mean-variance-normalized bimodality coefficient (BCmvn), and the expected number of doublets was adjusted for the estimated proportion of homotypic doublets. Singlets from the 15 samples were merged, log-normalized, scaled and reduced by principal component analysis. Batch effects were corrected by sample ID with Harmony (Korsunsky et al., 2019). Graph-based clustering (Louvain algorithm, resolution 2) was performed on the first 30 Harmony dimensions, and the cells were visualized by Uniform Manifold Approximation and Projection (UMAP). The main cell types were annotated from module scores (AddModuleScore, Seurat) of known signature markers (IGFBP6 for DSCs, PAEP for EEC, VWF for VEC, HBA1 for EB, CD68 for macrophages, HLA-DQA1 for cDC1, CLEC10A for cDC2, KLRC1 for NK cells, FCGR3B for neutrophils, MS4A2 for mast cells, CD3D for T cells and CD79A for B cells), following the annotation of Bao et al. (*45*). Clusters of low quality or ambiguous identity were excluded.

Neutrophil sub-clustering and differential expression: The neutrophil cluster was subset and re-clustered on the Harmony embedding (30 dimensions, resolution 2). The resulting subclusters were characterized by expression of CXCR4, CD63, ITGAM and MME, which correspond to the surface markers used for flow cytometry, and were assigned to three neutrophil states. Differential expression between neutrophils from RM patients and healthy donors was calculated with the FindMarkers function (Seurat) using a two-sided Wilcoxon rank-sum test. P < 0.05 was considered statistically significant.

### Cell invasion assay

To generate healthy neutrophil supernatants, PMNLs were isolated from the blood of healthy non-pregnant donors. 16,6 x 10^6^ cells were seeded in 1,5 mL RPMI -/- in a 6-well plate and cultured for 2 h. Cells were pelleted at 500 g for 5 min, and the supernatant was snap frozen. RPMI -/- was used in parallel as a negative control.

Invasion assays were performed using JEG-3 trophoblast cells (human choriocarcinoma-derived trophoblast cell line) as an established in vitro model to assess trophoblast migratory responses, as previously described (*31*). Briefly, 1 × 10^6^ JEG-3 cells were seeded in MEM+/+ in a 10 cm dish and grown for 48 h. Subsequently, cells were detached using Accutase and starved for 2h in MEM-/- to reduce basal proliferative activity and minimize interference from serum-derived invasion-promoting factors. A 24-well invasion chamber plate (#354480 Corning) was rehydrated with MEM-/- prior experiment for 2h. Upon starvation, cells were resuspended in a 1:10 dilution of neutrophil supernatants or 4nM NE (SE564, Elastin Products Company) in MEM-/- and incubated for 4 h. Control cells were treated with identically prepared RPMI-/- medium lacking neutrophil-derived factors. JEG-3 cells were subsequently washed with PBS-/-, counted and 50.000 cells/500µl MEM-/- were seeded per well in the upper chamber. The lower chamber was filled with 750µl MEM-/- supplemented with 10% FBS. After 24 h of invasion at 37°C, 5% CO2, cells were fixed and stained with Diff-Quick solution. Six images of each membrane were captured at 100× magnification using an Olympus BX53 microscope with an Olympus UC90 camera. CellSense Standard software was used for the analysis. ImageJ software (R Foundation) cell counter function was used to count the number of invaded cells.

### Statistical analysis

Statistical analyses were conducted using GraphPad Prism 9.1 (GraphPad Software) and R software (v4.4.2). Paired and unpaired t-tests were used to assess the statistically significant differences between two groups. Adjusted p-values were obtained using the Benjamini-Hochberg (BH) method. For comparisons involving more than two groups, one-way ANOVA followed by Tukey’s post-hoc test was applied. Linear regression analysis was used to investigate the relationship between the two variables. A significance threshold of P < 0.05 was considered statistically significant for all analyses.

## Supplementary Materials

Figs. S1 to S4 Tables S1 to S2

## Supporting information

Juvan_supplemental Files

## Acknowledgments

We are grateful to Sabine Kern for her excellent technical assistance.

## Funding

FWF grant P35294 (JK)

FFG-Bridge 1 grant 871284 (JK) OENB Anniversary Fund 17584 (JK)

Doctoral programs: FWF: DK-MOLIN (W1241) and DP-iDP (DOC-31) and OEAW (Doc Fellowship - 26477) (ZNJ, OK, ALS, NCM)

National Institutes of Health (NIH) R35GM137936 and P01HD106414 (BG)

Center for Human Systems Immunology at Stanford (BG), and the Stanford Maternal and Child Health Research Institute (BG, DF), Society for Reproductive Investigation Bayer Discovery/Innovation Grant (DF)

## Author contributions

Conceptualization: ZNJ, JK

Investigation: ZNJ, OK, TK, KM, NCM, EA, SR, ALS Formal Analysis: ZNJ, OK

Methodology: ZNJ, DF

Tissue pre-processing: ALH, SV, AL, DF Funding acquisition: JK, DF, BG Resources: JK, JP, MG, BG

Supervision: CW, JP, AH, GM, JF Visualization: ZNJ

Writing original draft: ZNJ, JK

Writing-review and editing: all co-authors

## Data and materials availability

All data are available in the main text or the supplementary materials.

---

**Fig. S1.**
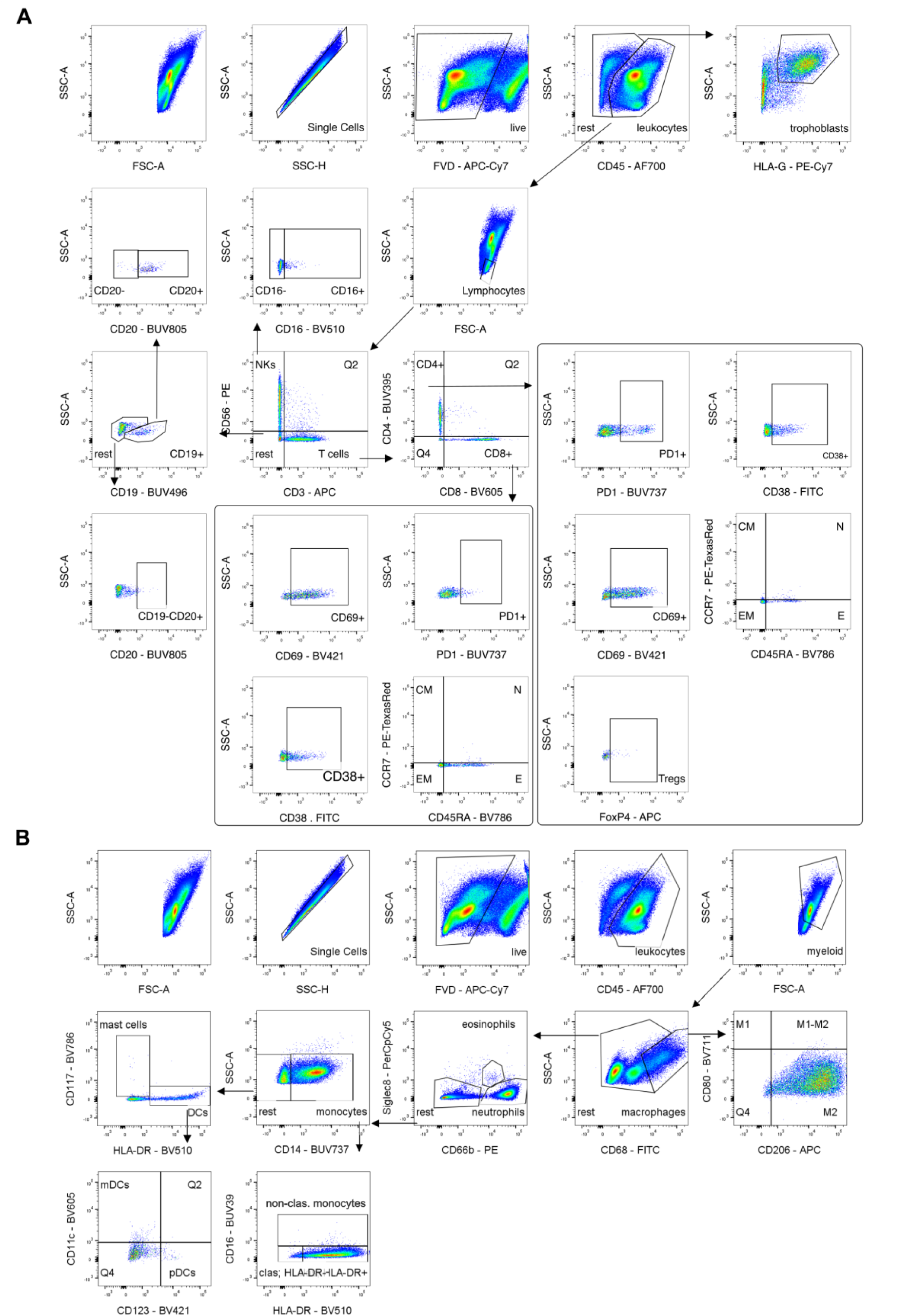
Gating strategy to determine immune populations in the tissue. (**A**) Doublets were removed using SSC-A and SSC-H, and live cells (FVD negative) were selected. CD45^+^ cells were gated to exclude non-immune cells, followed by a lymphocyte size gate. NK cells (CD56^+^, CD3^-^) and T cells (CD56^-^, CD3^+^) were separated. NK cells were split into cytotoxic (CD16^+^) and regulatory (CD16^-^) subsets. B cells were identified as CD3^-^, CD56^-^, CD19^+/-^, CD20^+/-^. T cells were divided into CD4^+^ and CD8^+^ subsets, and their activation was assessed using PD-1, CD38, and CD69. CD4^+^ and CD8^+^ cells were further classified into CM, N, E, and EM subpopulations based on CCR7 and CD45RA expression. Tregs were identified as FoxP3^+^ within CD4^+^ cells. (**B**) Gates for single cells, live cells, and leukocytes were applied, followed by a myeloid gate. Macrophages (CD68high) were subdivided into M1 (CD80^+^, CD206^-^), M1-M2 (CD80^+^, CD206^+^), and M2 (CD80^-^, CD206^+^). Eosinophils were Siglec8^+^, and neutrophils were CD66b^+^, Siglec8^-^. Monocytes (CD14^+^) were split into non-classical (CD16^+^) and classical (CD16^-^), with HLA-DR assessed in classical monocytes. CD14^-^ cells were identified as mast cells (CD117^+^, HLA-DR^-^) and dendritic cells (CD117^-^, HLA-DR^+^). DCs were further categorized as mDCs (CD11c^+^, CD123^-^) and pDCs (CD11c^-^, CD123^+^). Activation of myeloid populations was evaluated based on PD-L1 expression.

**Fig. S2.**
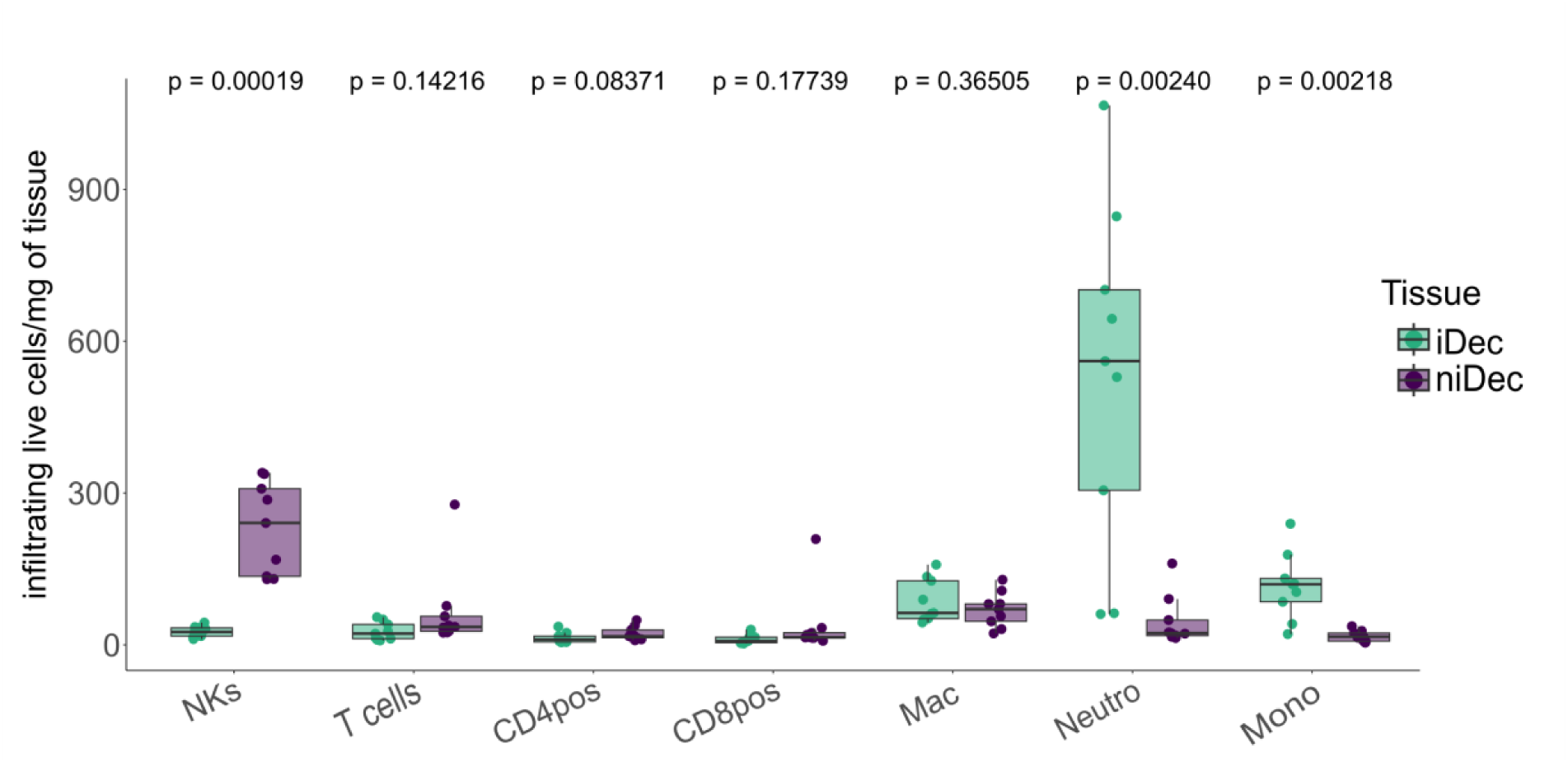
Number of immune cells present in the tissue determined by flow cytometry. Average of main immune populations in iDec and niDec represented as live cells/mg of tissue (N=9). Paired Student’s t-tests were conducted for comparing two groups. Data are presented as mean ± SD, *P < 0.05.

**Fig. S3.**
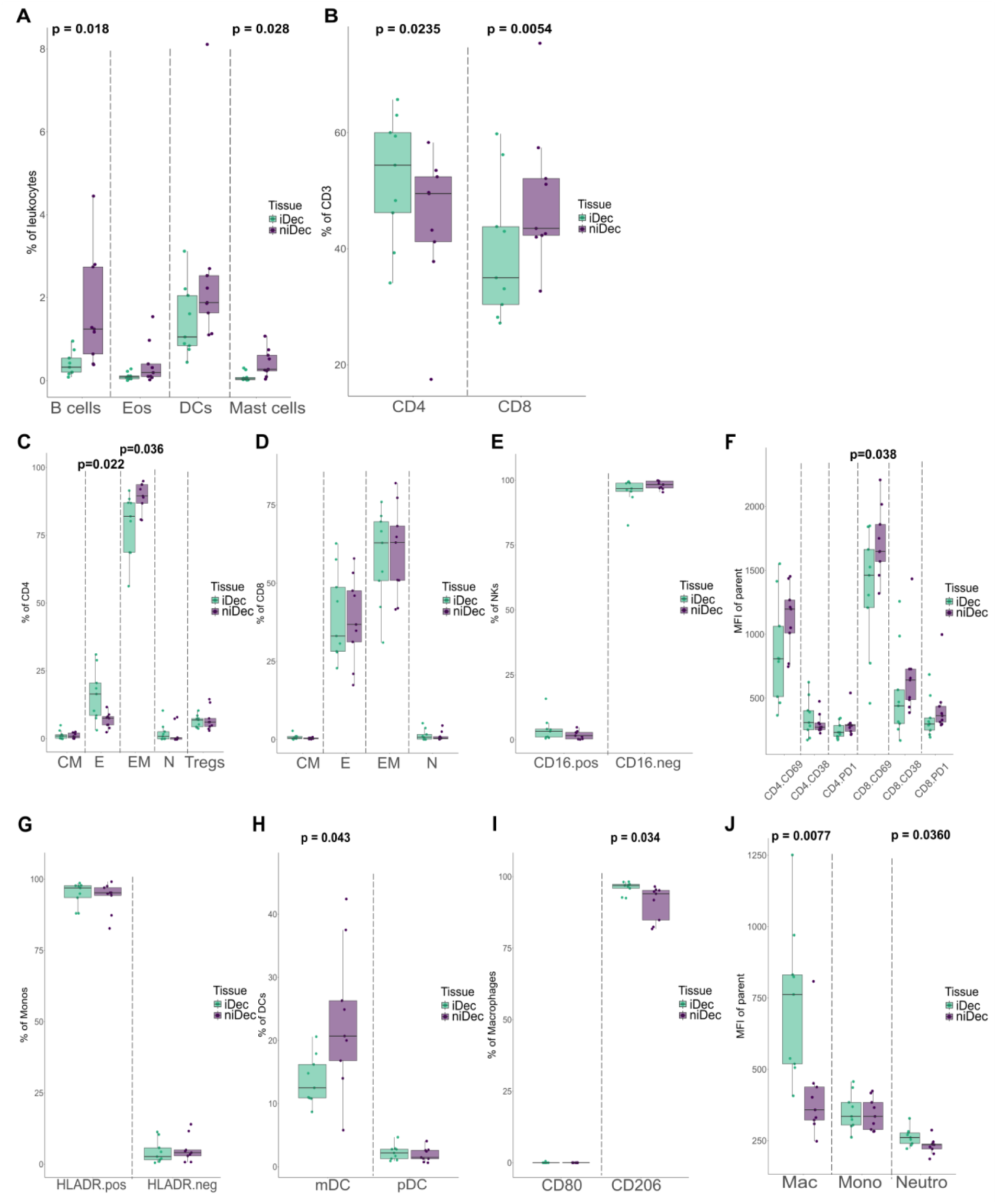
Immune microenvironment in iDec and niDec. (**A**) Immune subpopulations in iDec and niDec represented as a percentage of leukocytes (N=9). (**B**) Presence of CD4^+^ and CD8^+^ Tcells (CD3^+^) in iDec and niDec represented as a percentage of CD3^+^ cells (N=9). (**C**) Distribution of CM (CCR7^+^, CD45RA^-^), N (CCR7^+^, CD45RA^+^), E (CCR7^-^, CD45RA^+^), and EM (CCR7^-^, CD45RA^-^) subsets of CD4^+^ cells. Additionally, Tregs were identified as a subgroup. (D) Distribution of CM (CCR7^+^, CD45RA^-^), N (CCR7^+^, CD45RA^+^), E (CCR7^-^, CD45RA^+^), and EM (CCR7^-^, CD45RA^-^) subsets of CD8^+^ cells. (**E**) Cytotoxic (CD16^+^) and regulatory (CD16^-^) subsets of NKs displayed as a percentage of NKs. (**F**) Activation status of CD4^+^ and CD8^+^ cells based on the expression of PD-1, CD38, and CD69, presented as MFI. (**G**) HLA-DR expression on monocytes. (**H**) Distribution of DC subsets, including mDCs (CD11b^+^) and pDCs (CD123^+^) cells. (**I**) Macrophage subsets categorized as M1 (CD80^+^) and M2 (CD206^+^), presented as a percentage of macrophages. (**J**) PD-L1 expression on larger myeloid populations, such as macrophages, monocytes, and neutrophils. Paired Student’s t-tests were conducted for comparing two groups. Data are presented as mean ± SD, *P < 0.05.

**Fig. S4.**
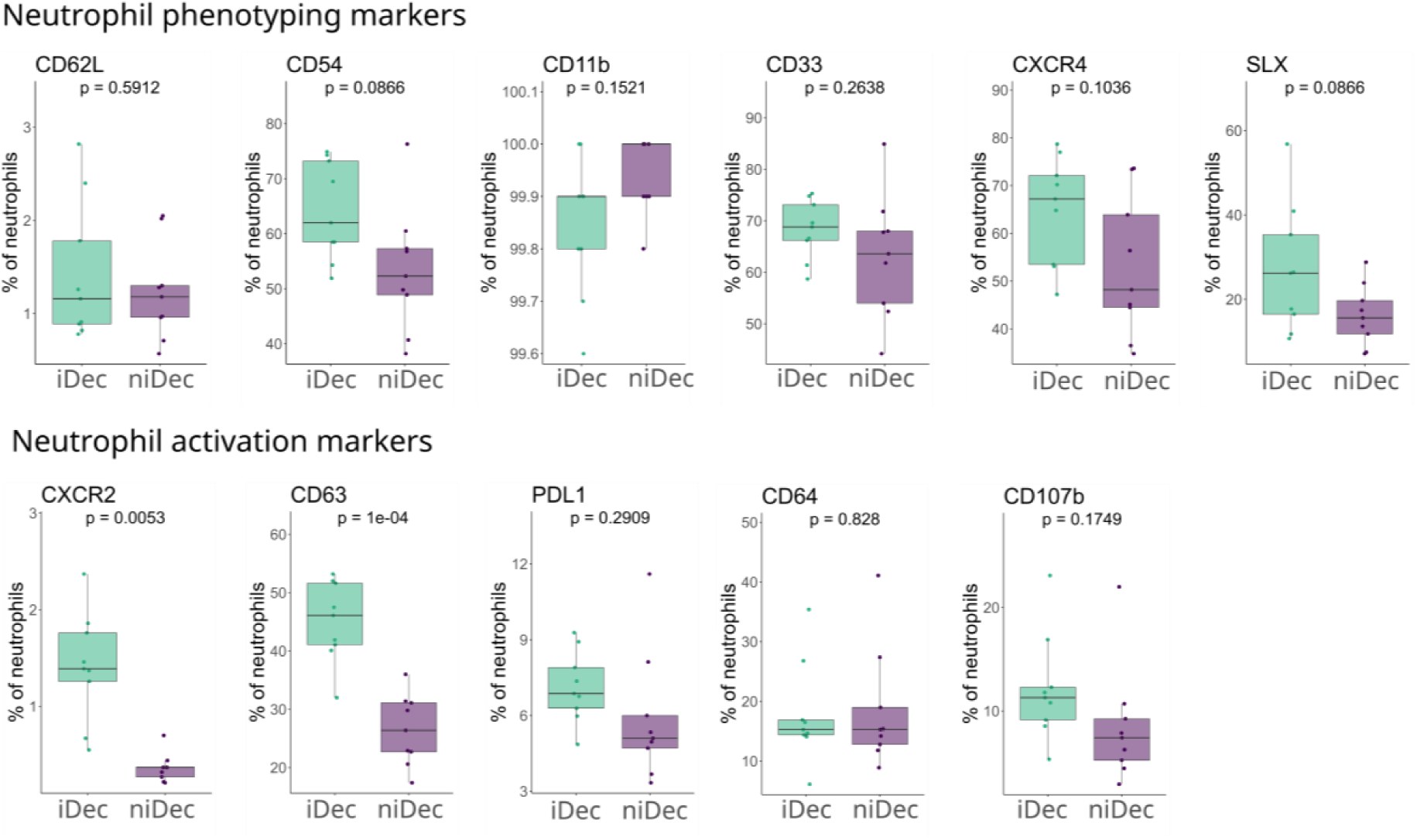
Neutrophil marker expression in iDec and niDec. Expression of known neutrophil phenotyping and activation markers. Paired Student’s t-tests were conducted for comparing two groups. Data are presented as mean ± SD, *P < 0.05.

**Table S1.**
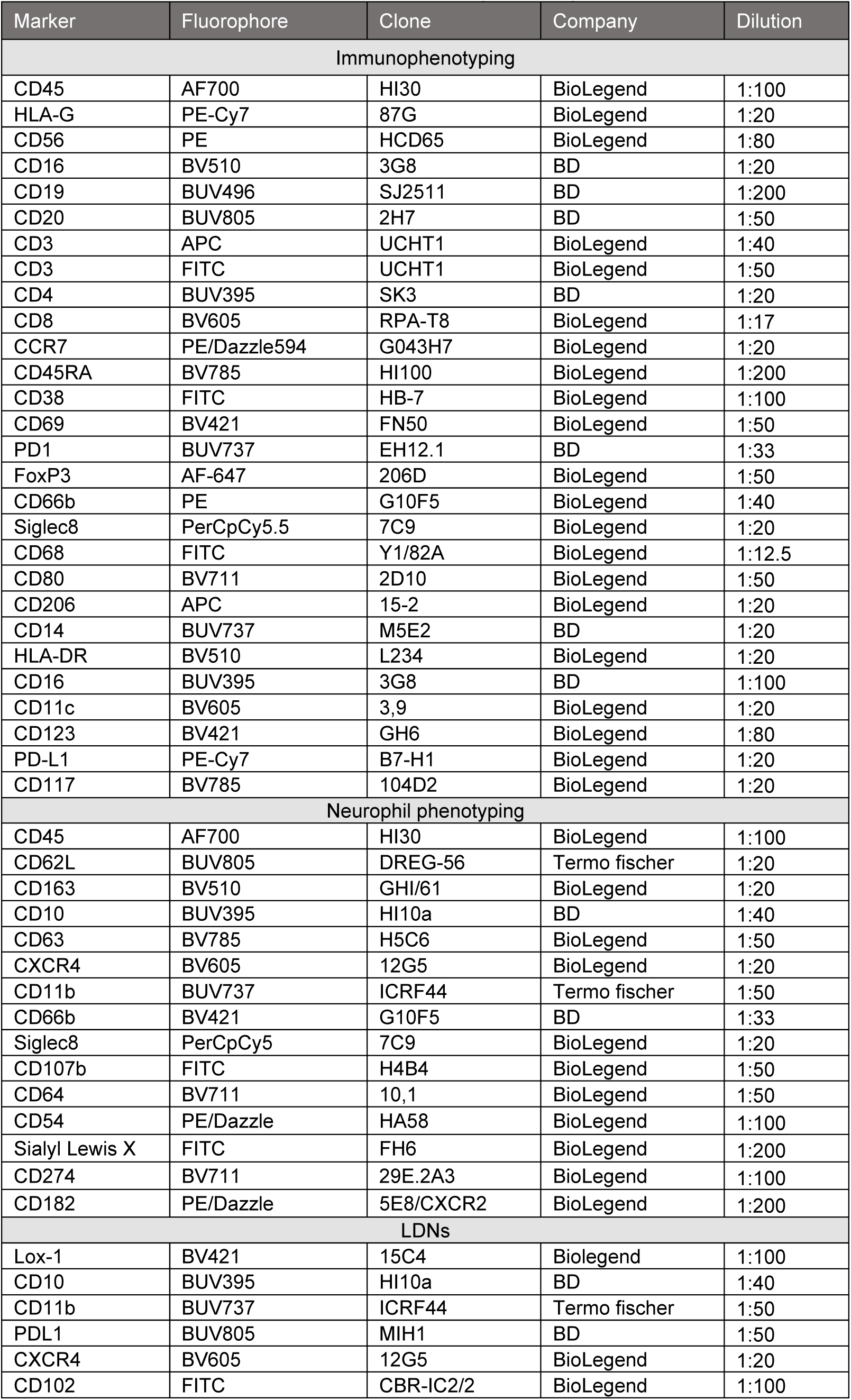
List of antibodies used for flow cytometry.

**Table S2:**
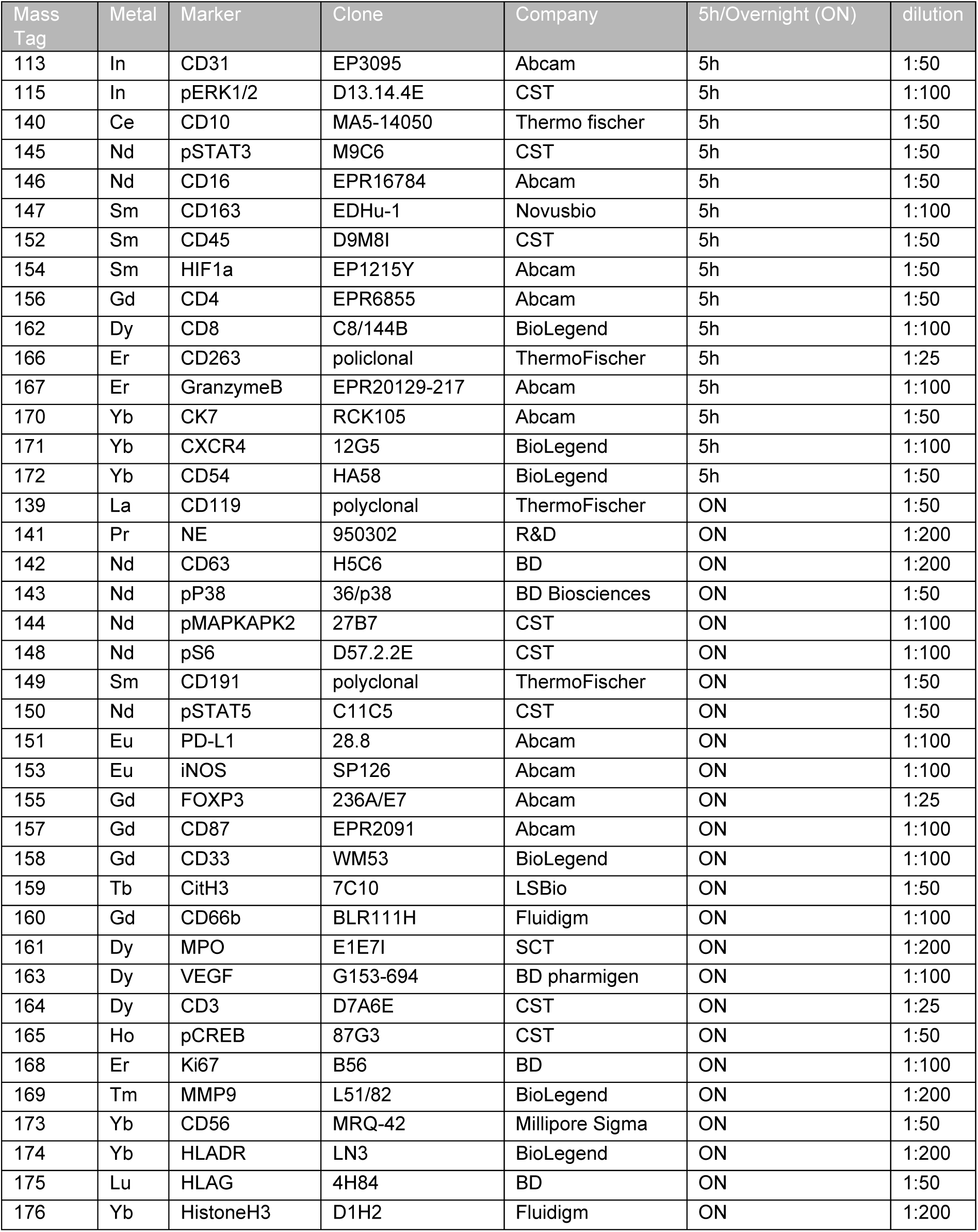
List of antibodies used for IMC.

## REFERENCES AND NOTES

1. Knöfler M, Haider S, Saleh L, Pollheimer J, Gamage TKJB, James J. Human placenta and trophoblast development: key molecular mechanisms and model systems. Cell Mol Life Sci. 2019 Sep;76(18):3479–96. doi:10.1007/s00018-019-03104-6

2. 0.000002. Turco MY, Moffett A. Development of the human placenta. Development. 2019 Nov 15;146(22):dev163428. doi:10.1242/dev.163428

3. Ng SW, Norwitz GA, Pavlicev M, Tilburgs T, Simón C, Norwitz ER. Endometrial Decidualization: The Primary Driver of Pregnancy Health. Int J Mol Sci. 2020 Jun 8;21(11):4092. doi:10.3390/ijms21114092

4. Rana S, Lemoine E, Granger JP, Karumanchi SA. Preeclampsia: Pathophysiology, Challenges, and Perspectives. Circ Res. 2019 Mar 29;124(7):1094–112. doi:10.1161/CIRCRESAHA.118.313276

5. Lin RC, Chao YY, Su MT, Tsai HL, Tsai PY, Wang CY. Upregulation of miR-20b-5p inhibits trophoblast invasion by blocking autophagy in recurrent miscarriage. Cell Signal. 2024 Jan;113:110934. doi:10.1016/j.cellsig.2023.110934

6. Suryawanshi H, Morozov P, Straus A, Sahasrabudhe N, Max KEA, Garzia A, et al. A single-cell survey of the human first-trimester placenta and decidua. Sci Adv. 2018 Oct 5;4(10):eaau4788. doi:10.1126/sciadv.aau4788

7. Guo C, Cai P, Jin L, Sha Q, Yu Q, Zhang W, et al. Single-cell profiling of the human decidual immune microenvironment in patients with recurrent pregnancy loss. Cell Discov. 2021 Jan 4;7(1):1. doi:10.1038/s41421-020-00236-z

8. Hou R, Huang R, Zhou Y, Lin D, Xu J, Yang L, et al. Single-cell profiling of the microenvironment in decidual tissue from women with missed abortions. Fertil Steril. 2023 Mar;119(3):492–503. doi:10.1016/j.fertnstert.2022.12.016

9. Moser G, Kummer D, Gruber M, Hartmann T, Forstner D, Guettler J, et al. Remodeling lesions locate at sites of strong extravillous trophoblast invasion and are associated with neutrophil presence in the human first-trimester decidua. Hum Reprod. 2026 Jul 1;41(7):1078–96. doi:10.1093/humrep/deag078

10. Höbler AL, Lackner AI, Öhler B, Musiejovsky L, Haslinger P, Vondra S, et al. Fibrinoid-associated neutrophils promote invasive placentation and neutrophil reprogramming at the maternal-fetal interface. Sci Immunol. 2026 Sep 25;11(123):eaea4118. doi:10.1126/sciimmunol.aea4118

11. Rosales C. Neutrophils at the crossroads of innate and adaptive immunity. J Leukoc Biol. 2020 Jul;108(1):377–96. doi:10.1002/JLB.4MIR0220-574RR PubMed PMID: 32202340.

12. Rosales C. Neutrophil: A Cell with Many Roles in Inflammation or Several Cell Types? Front Physiol. 2018 Feb 20;9:113. doi:10.3389/fphys.2018.00113

13. Ng LG, Ostuni R, Hidalgo A. Heterogeneity of neutrophils. Nat Rev Immunol. 2019 Apr;19(4):255–65. doi:10.1038/s41577-019-0141-8 PubMed PMID: 30816340.

14. Hellebrekers P, Vrisekoop N, Koenderman L. Neutrophil phenotypes in health and disease. Eur J Clin Invest. 2018 Nov;48:e12943. doi:10.1111/eci.12943

15. Nadkarni S, Smith J, Sferruzzi-Perri AN, Ledwozyw A, Kishore M, Haas R, et al. Neutrophils induce proangiogenic T cells with a regulatory phenotype in pregnancy. Proc Natl Acad Sci U S A. 2016 Dec 27;113(52):E8415–24. doi:10.1073/pnas.1611944114 PubMed PMID: 27956610;

16. PubMed Central PMCID: PMC5206541.

16. Ostrand-Rosenberg S, Sinha P, Figley C, Long R, Park D, Carter D, et al. Frontline Science: Myeloid-derived suppressor cells (MDSCs) facilitate maternal-fetal tolerance in mice. J Leukoc Biol. 2017 May;101(5):1091–101. doi:10.1189/jlb.1HI1016-306RR PubMed PMID: 28007981; PubMed Central PMCID: PMC5380379.

17. Dockree S, Shine B, Pavord S, Impey L, Vatish M. White blood cells in pregnancy: reference intervals for before and after delivery. eBioMedicine. 2021 Dec;74:103715. doi:10.1016/j.ebiom.2021.103715

18. Carlucci PM, Purmalek MM, Dey AK, Temesgen-Oyelakin Y, Sakhardande S, Joshi AA, et al. Neutrophil subsets and their gene signature associate with vascular inflammation and coronary atherosclerosis in lupus. JCI Insight. 2018 Apr 19;3(8):e99276. doi:10.1172/jci.insight.99276

19. Valadez-Cosmes P, Maitz K, Kindler O, Raftopoulou S, Kienzl M, Santiso A, et al. Identification of Novel Low-Density Neutrophil Markers Through Unbiased High-Dimensional Flow Cytometry Screening in Non-Small Cell Lung Cancer Patients. Front Immunol. 2021 Aug 13;12:703846. doi:10.3389/fimmu.2021.703846

20. Ye H, Li L, Dong Y, Zheng Q, Sha Y, Li L, et al. Dysregulated low-density granulocyte contributes to early spontaneous abortion. Front Immunol. 2023 Feb 23;14:1119756. doi:10.3389/fimmu.2023.1119756

21. Grieshaber-Bouyer R, Radtke FA, Cunin P, Stifano G, Levescot A, Vijaykumar B, et al. The neutrotime transcriptional signature defines a single continuum of neutrophils across biological compartments. Nat Commun. 2021 May 17;12(1):2856. doi:10.1038/s41467-021-22973-9

22. Price TH, Dale DC. Neutrophil preservation: the effect of short-term storage on in vivo kinetics. J Clin Invest. 1977 Mar;59(3):475–80. doi:10.1172/JCI108662 PubMed PMID: 838860; PubMed Central PMCID: PMC333384.

23. McKenna E, Mhaonaigh AU, Wubben R, Dwivedi A, Hurley T, Kelly LA, et al. Neutrophils: Need for Standardized Nomenclature. Front Immunol. 2021 Apr 15;12:602963. doi:10.3389/fimmu.2021.602963

24. Kargl J, Busch SE, Yang GHY, Kim KH, Hanke ML, Metz HE, et al. Neutrophils dominate the immune cell composition in non-small cell lung cancer. Nat Commun. 2017 Feb 1;8:14381. doi:10.1038/ncomms14381 PubMed PMID: 28146145; PubMed Central PMCID: PMC5296654.

25. Solders M, Gorchs L, Gidlöf S, Tiblad E, Lundell AC, Kaipe H. Maternal Adaptive Immune Cells in Decidua Parietalis Display a More Activated and Coinhibitory Phenotype Compared to Decidua Basalis. Stem Cells Int. 2017;2017:8010961. doi:10.1155/2017/8010961 PubMed PMID: 29317870; PubMed Central PMCID: PMC5727765.

26. Vondra S, Höbler AL, Lackner AI, Raffetseder J, Mihalic ZN, Vogel A, et al. The human placenta shapes the phenotype of decidual macrophages. Cell Rep. 2023 Jan 10;42(1):111977. doi:10.1016/j.celrep.2022.111977 PubMed PMID: 36640334.

27. Moser G, Orendi K, Gauster M, Siwetz M, Helige C, Huppertz B. The art of identification of extravillous trophoblast. Placenta. 2011 Feb;32(2):197–9. doi:10.1016/j.placenta.2010.11.008 PubMed PMID: 21183217.

28. Mattos MS, Lopes ME, De Araujo AM, Alvarenga DM, Nakagaki BN, Mafra K, et al. Prolonged neutrophil survival at necrotic sites is a fundamental feature for tissue recovery and resolution of hepatic inflammation. J Leukoc Biol. 2020 Oct 1;108(4):1199–213. doi:10.1002/JLB.1MA0420-634R

29. Vento-Tormo R, Efremova M, Botting RA, Turco MY, Vento-Tormo M, Meyer KB, et al. Single-cell reconstruction of the early maternal–fetal interface in humans. Nature. 2018 Nov 15;563(7731):347–53. doi:10.1038/s41586-018-0698-6

30. Othman A, Sekheri M, Filep JG. Roles of neutrophil granule proteins in orchestrating inflammation and immunity. FEBS J. 2022 Jul;289(14):3932–53. doi:10.1111/febs.15803

31. Mihalic ZN, Kloimböck T, Cosic-Mujkanovic N, Valadez-Cosmes P, Maitz K, Kindler O, et al. Myeloperoxidase enhances the migration and invasion of human choriocarcinoma JEG-3 cells. Redox Biol. 2023 Nov;67:102885. doi:10.1016/j.redox.2023.102885 PubMed PMID: 37776707; PubMed Central PMCID: PMC10556814.

32. Croxatto D, Micheletti A, Montaldo E, Orecchia P, Loiacono F, Canegallo F, et al. Group 3 innate lymphoid cells regulate neutrophil migration and function in human decidua. Mucosal Immunol. 2016 Nov;9(6):1372–83. doi:10.1038/mi.2016.10 PubMed PMID: 26906405.

33. Adrover JM, Del Fresno C, Crainiciuc G, Cuartero MI, Casanova-Acebes M, Weiss LA, et al. A Neutrophil Timer Coordinates Immune Defense and Vascular Protection. Immunity. 2019 Feb;50(2):390–402.e10. doi:10.1016/j.immuni.2019.01.002

34. Zhang X, Paré P, Sandford A. PMN degranulation in relation to CD63 expression and genetic polymorphisms in healthy individuals and COPD patients. Int J Mol Med. 2007 May 1. doi:10.3892/ijmm.19.5.817

35. Kuhns DB, Long Priel DA, Gallin JI. Loss of L-selectin (CD62L) on human neutrophils following exudation in vivo. Cell Immunol. 1995 Sep;164(2):306–10. doi:10.1006/cimm.1995.1174 PubMed PMID: 7544694.

36. Amsalem H, Kwan M, Hazan A, Zhang J, Jones RL, Whittle W, et al. Identification of a novel neutrophil population: proangiogenic granulocytes in second-trimester human decidua. J Immunol Baltim Md 1950. 2014 Sep 15;193(6):3070–9. doi:10.4049/jimmunol.1303117 PubMed PMID: 25135830.

37. Parthasarathy U, Kuang Y, Thakur G, Hogan JD, Wyche TP, Norton JE, et al. Distinct subsets of neutrophils crosstalk with cytokines and metabolites in patients with sepsis. iScience. 2023 Feb 17;26(2):105948. doi:10.1016/j.isci.2023.105948 PubMed PMID: 36756375; PubMed Central PMCID: PMC9900520.

38. Sun R, Huang J, Yang Y, Liu L, Shao Y, Li L, et al. Dysfunction of low-density neutrophils in peripheral circulation in patients with sepsis. Sci Rep. 2022 Jan 13;12(1):685. doi:10.1038/s41598-021-04682-x

39. Condamine T, Dominguez GA, Youn JI, Kossenkov AV, Mony S, Alicea-Torres K, et al. Lectin-type oxidized LDL receptor-1 distinguishes population of human polymorphonuclear myeloid-derived suppressor cells in cancer patients. Sci Immunol. 2016 Aug 5;1(2). doi:10.1126/sciimmunol.aaf8943

40. Yang W, Tao Y, Wu Y, Zhao X, Ye W, Zhao D, et al. Neutrophils promote the development of reparative macrophages mediated by ROS to orchestrate liver repair. Nat Commun. 2019 Mar 6;10(1):1076. doi:10.1038/s41467-019-09046-8

41. Jaeger BN, Donadieu J, Cognet C, Bernat C, Ordoñez-Rueda D, Barlogis V, et al. Neutrophil depletion impairs natural killer cell maturation, function, and homeostasis. J Exp Med. 2012 Mar 12;209(3):565–80. doi:10.1084/jem.20111908

42. Zhang Y, Aldo P, You Y, Ding J, Kaislasuo J, Petersen JF, et al. Trophoblast-secreted soluble-PD-L1 modulates macrophage polarization and function. J Leukoc Biol. 2020 Sep;108(3):983–98. doi:10.1002/JLB.1A0420-012RR

43. Tilburgs T, Evans JH, Crespo ÂC, Strominger JL. The HLA-G cycle provides for both NK tolerance and immunity at the maternal–fetal interface. Proc Natl Acad Sci. 2015 Oct 27;112(43):13312–7. doi:10.1073/pnas.1517724112

44. Wienke J, Brouwers L, van der Burg LM, Mokry M, Scholman RC, Nikkels PGJ, et al. Human Tregs at the materno-fetal interface show site-specific adaptation reminiscent of tumor Tregs. JCI Insight. 2020 Sep 17;5(18):e137926. doi:10.1172/jci.insight.137926

45. Bao S, Chen Z, Qin D, Xu H, Deng X, Zhang R, et al. Single-cell profiling reveals mechanisms of uncontrolled inflammation and glycolysis in decidual stromal cell subtypes in recurrent miscarriage. Hum Reprod. 2023 Jan 5;38(1):57–74. doi:10.1093/humrep/deac240

46. Greenbaum S, Averbukh I, Soon E, Rizzuto G, Baranski A, Greenwald NF, et al. A spatially resolved timeline of the human maternal–fetal interface. Nature. 2023 Jul 20;619(7970):595–605. doi:10.1038/s41586-023-06298-9

47. Krop J, van der Zwan A, Ijsselsteijn ME, Kapsenberg H, Luk SJ, Hendriks SH, et al. Imaging mass cytometry reveals the prominent role of myeloid cells at the maternal-fetal interface. iScience. 2022 Jul 15;25(7):104648. doi:10.1016/j.isci.2022.104648 PubMed PMID: 35811852; PubMed Central PMCID: PMC9257341.

48. Hebeda CB, Savioli AC, Scharf P, de Paula-Silva M, Gil CD, Farsky SHP, et al. Neutrophil depletion in the pre-implantation phase impairs pregnancy index, placenta and fetus development. Front Immunol. 2022 Sep 29;13:969336. doi:10.3389/fimmu.2022.969336

49. Wickham H, Averick M, Bryan J, Chang W, McGowan LD, François R, Wickham, et al., (2019). Welcome to the Tidyverse. Journal of Open Source Software, 4(43), 1686.

50. Stelzer IA, Ghaemi MS, Han X, Ando K, Hédou JJ, Feyaerts D, et al. Integrated trajectories of the maternal metabolome, proteome, and immunome predict labor onset. Sci Transl Med. 2021 May 5;13(592):eabd9898. doi:10.1126/scitranslmed.abd9898 PubMed PMID: 33952678; PubMed Central PMCID: PMC8136601.

51. Greenwald NF, Miller G, Moen E, Kong A, Kagel A, Dougherty T, et al. Whole-cell segmentation of tissue images with human-level performance using large-scale data annotation and deep learning. Nat Biotechnol. 2022 Apr;40(4):555–65. doi:10.1038/s41587-021-01094-0

52. Windhager J, Bodenmiller B, Eling N. An end-to-end workflow for multiplexed image processing and analysis [preprint] [Internet]. Bioinformatics; 2021 Nov [cited 2023 Sep 18]. Available from: http://biorxiv.org/lookup/doi/10.1101/2021.11.12.468357 doi:10.1101/2021.11.12.468357

