## Supplementary material for "The decidual microenvironment shapes neutrophil subsets that promote extravillous trophoblast invasion in early pregnancy": Juvan_supplemental Files

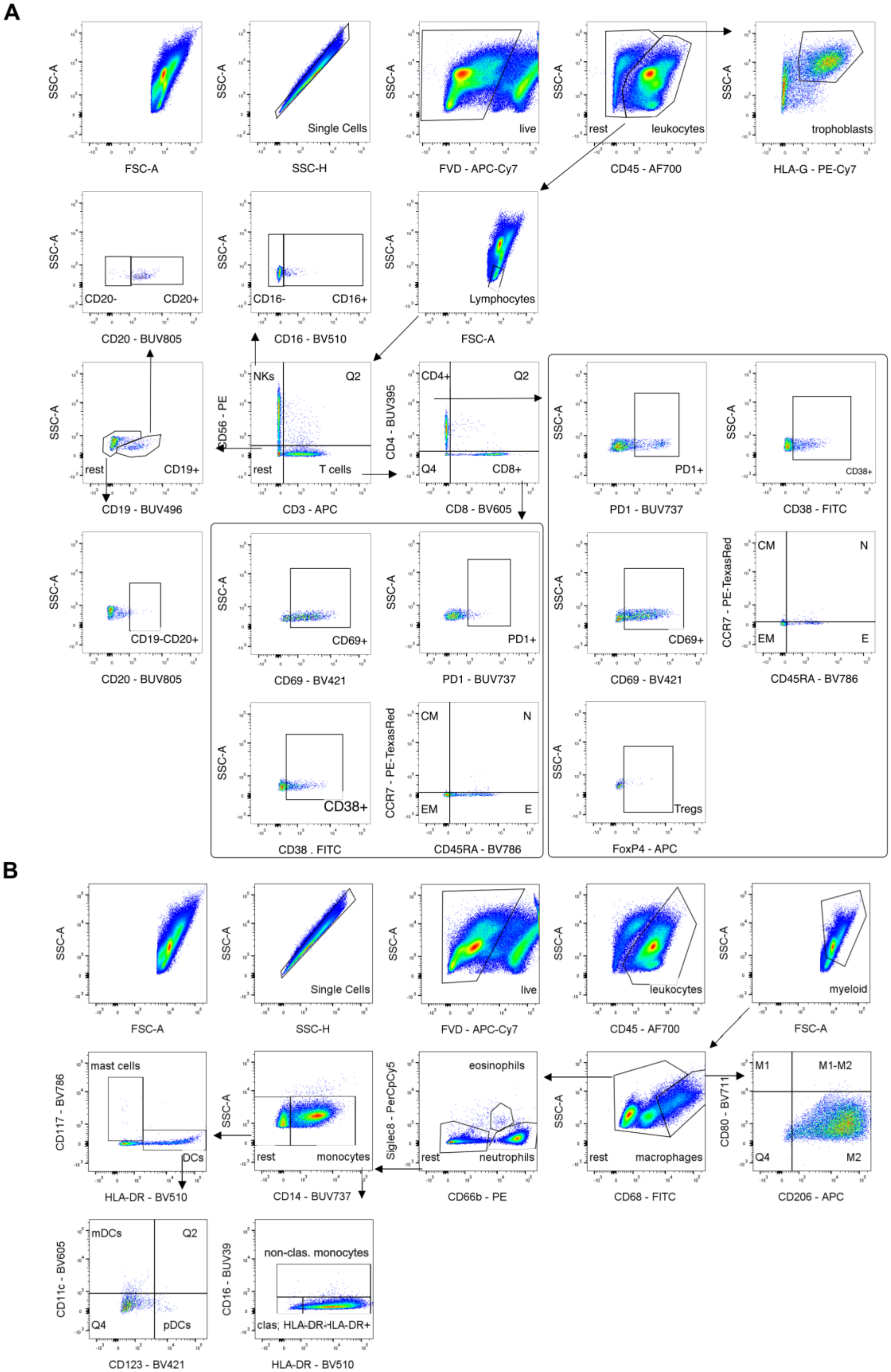
A
SSC-A
FSC-A
SSC-H
Single Cells
FVD - APC-Cy7
live
CD45 - AF700
rest
leukocytes
HLA-G - PE-Cy7
trophoblasts
CD20 - BUV805
CD20-
CD20+
CD16 - BV510
CD16-
CD16+
Lymphocytes
CD19 - BUV496
CD19+
CD3 - APC
CD56 - PE
NKs
Q2
T cells
CD4 - BUV395
CD8 - BV605
CD4+
CD8+
Q4
PD1 - BUV737
PD1+
CD38 - FITC
CD38+
CD69 - BV421
CD69+
CD19-CD20+
CCR7 - PE-TexasRed
CD45RA - BV786
CM
N
EM
E
CD38 . FITC
FoxP4 - APC
Tregs
B
myeloid
CD117 - BV786
mast cells
HLA-DR - BV510
DCs
CD14 - BUV737
monocytes
Siglec8 - PerCpCy5
CD66b - PE
eosinophils
neutrophils
CD68 - FITC
macrophages
CD80 - BV711
CD206 - APC
M1
M1-M2
M2
CD11c - BV605
CD123 - BV421
mDCs
pDCs
CD16 - BUV39
non-clas. monocytes
clas; HLA-DR-HLA-DR+

**Fig. S1. Gating strategy to determine immune populations in the tissue.**
(**A**) Doublets were removed using SSC-A and SSC-H, and live cells (FVD negative) were selected. $CD45^+$ cells were gated to exclude non-immune cells, followed by a lymphocyte size gate. NK cells ($CD56^+$, $CD3^-$) and T cells ($CD56^-$, $CD3^+$) were separated. NK cells were split into cytotoxic ($CD16^+$) and regulatory ($CD16^-$) subsets. B cells were identified as $CD3^-$, $CD56^-$, $CD19^{+/-}$, $CD20^{+/-}$. T cells were divided into $CD4^+$ and $CD8^+$ subsets, and their activation was assessed using PD-1, CD38, and CD69. $CD4^+$ and $CD8^+$ cells were further classified into CM, N, E, and EM subpopulations based on CCR7 and CD45RA expression. Tregs were identified as $FoxP3^+$ within $CD4^+$ cells. (**B**) Gates for single cells, live cells, and leukocytes were applied, followed by a myeloid gate. Macrophages (CD68high) were subdivided into M1 ($CD80^+$, $CD206^-$), M1-M2 ($CD80^+$, $CD206^+$), and M2 ($CD80^-$, $CD206^+$). Eosinophils were $Siglec8^+$, and neutrophils were $CD66b^+$, $Siglec8^-$. Monocytes ($CD14^+$) were split into non-classical ($CD16^+$) and classical ($CD16^-$), with HLA-DR assessed in classical monocytes. $CD14^-$ cells were identified as mast cells ($CD117^+$, $HLA\text{-}DR^-$) and dendritic cells ($CD117^-$, $HLA\text{-}DR^+$). DCs were further categorized as mDCs ($CD11c^+$, $CD123^-$) and pDCs ($CD11c^-$, $CD123^+$). Activation of myeloid populations was evaluated based on PD-L1 expression.

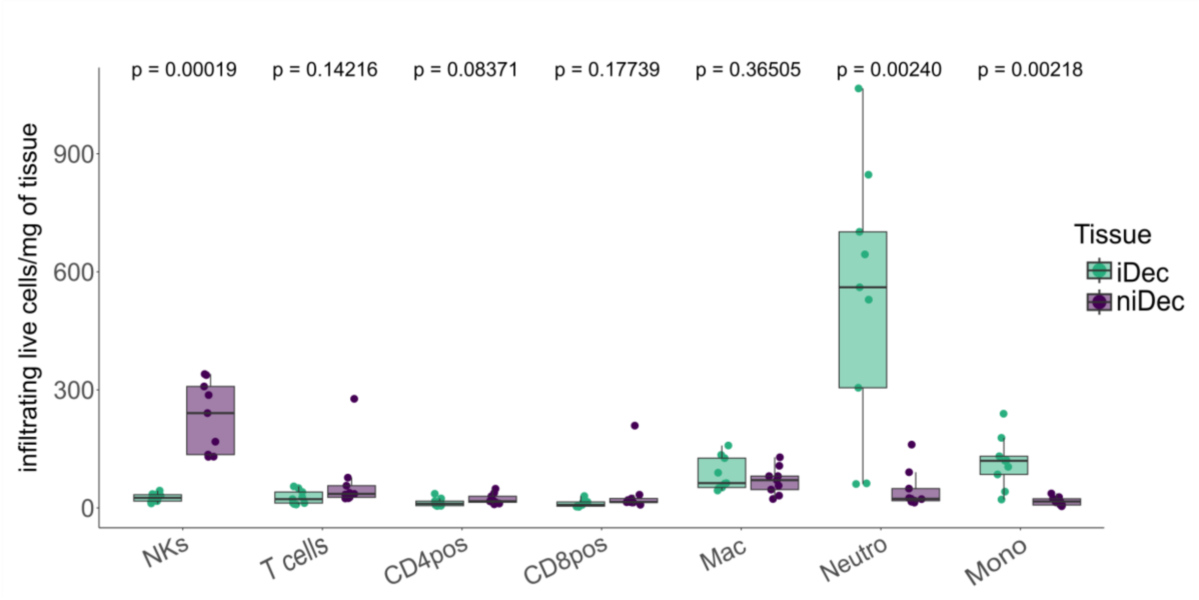


**Fig. S2. Number of immune cells present in the tissue determined by flow cytometry.**
Average of main immune populations in iDec and niDec represented as live cells/mg of tissue (N=9). Paired Student's t-tests were conducted for comparing two groups. Data are presented as mean ± SD, $*P < 0.05$.

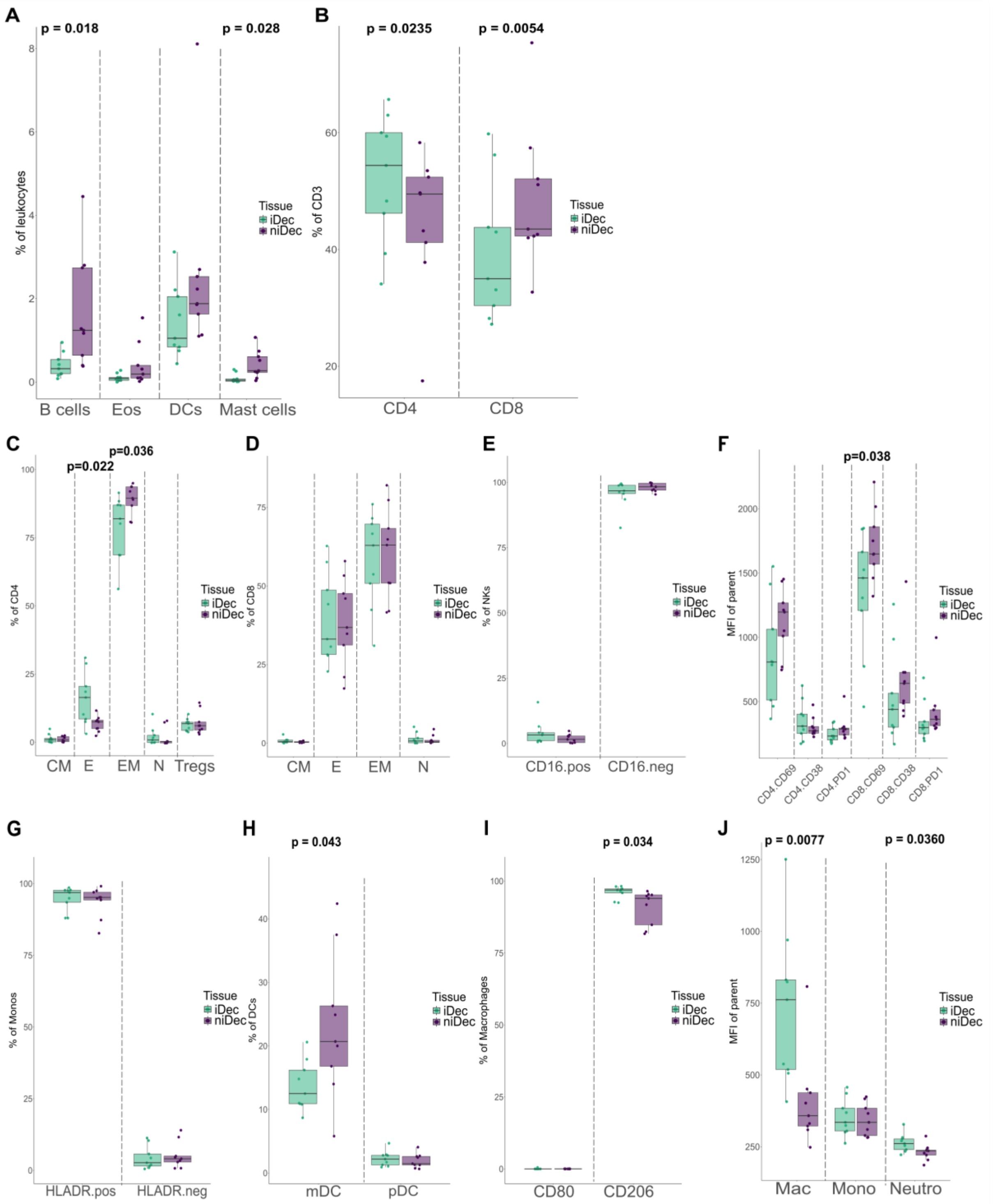


**Fig. S3. Immune microenvironment in iDec and niDec.**
(**A**) Immune subpopulations in iDec and niDec represented as a percentage of leukocytes (N=9). (**B**) Presence of $CD4^+$ and $CD8^+$ Tcells ($CD3^+$) in iDec and niDec represented as a percentage of $CD3^+$ cells (N=9). (**C**) Distribution of CM ($CCR7^+$, $CD45RA^-$), N ($CCR7^+$, $CD45RA^+$), E ($CCR7^-$, $CD45RA^+$), and EM ($CCR7^-$, $CD45RA^-$) subsets of $CD4^+$ cells. Additionally, Tregs were identified as a subgroup. (D) Distribution of CM ($CCR7^+$, $CD45RA^-$), N ($CCR7^+$, $CD45RA^+$), E

(CCR7$^-$, CD45RA$^+$), and EM (CCR7$^-$, CD45RA$^-$) subsets of CD8$^+$ cells. (**E**) Cytotoxic (CD16$^+$) and regulatory (CD16$^-$) subsets of NKs displayed as a percentage of NKs. (**F**) Activation status of CD4$^+$ and CD8$^+$ cells based on the expression of PD-1, CD38, and CD69, presented as MFI. (**G**) HLA-DR expression on monocytes. (**H**) Distribution of DC subsets, including mDCs (CD11b$^+$) and pDCs (CD123$^+$) cells. (**I**) Macrophage subsets categorized as M1 (CD80$^+$) and M2 (CD206$^+$), presented as a percentage of macrophages. (**J**) PD-L1 expression on larger myeloid populations, such as macrophages, monocytes, and neutrophils. Paired Student's t-tests were conducted for comparing two groups. Data are presented as mean ± SD, *$P < 0.05$.

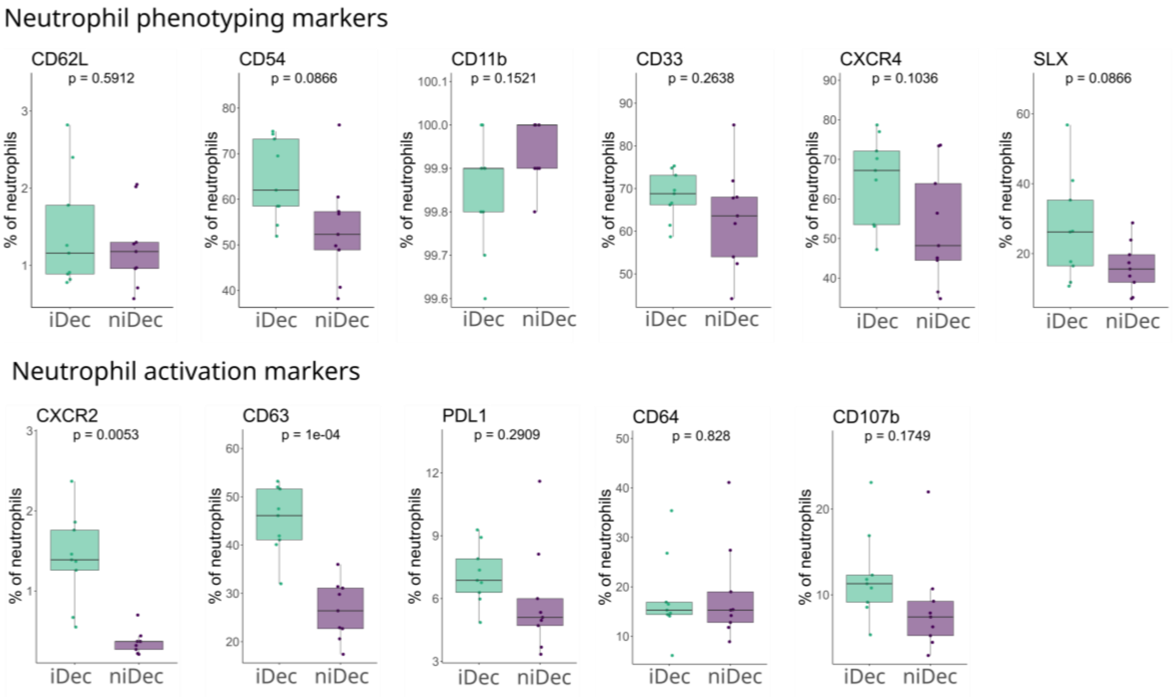


**Fig. S4. Neutrophil marker expression in iDec and niDec.**
Expression of known neutrophil phenotyping and activation markers. Paired Student's t-tests were conducted for comparing two groups. Data are presented as mean ± SD, *$P < 0.05$.

**Table S1. List of antibodies used for flow cytometry.**

| Marker | Fluorophore | Clone | Company | Dilution |
|---|---|---|---|---|
| Immunophenotyping | | | | |
| CD45 | AF700 | HI30 | BioLegend | 1:100 |
| HLA-G | PE-Cy7 | 87G | BioLegend | 1:20 |
| CD56 | PE | HCD65 | BioLegend | 1:80 |
| CD16 | BV510 | 3G8 | BD | 1:20 |
| CD19 | BUV496 | SJ2511 | BD | 1:200 |
| CD20 | BUV805 | 2H7 | BD | 1:50 |
| CD3 | APC | UCHT1 | BioLegend | 1:40 |
| CD3 | FITC | UCHT1 | BioLegend | 1:50 |
| CD4 | BUV395 | SK3 | BD | 1:20 |
| CD8 | BV605 | RPA-T8 | BioLegend | 1:17 |
| CCR7 | PE/Dazzle594 | G043H7 | BioLegend | 1:20 |
| CD45RA | BV785 | HI100 | BioLegend | 1:200 |
| CD38 | FITC | HB-7 | BioLegend | 1:100 |
| CD69 | BV421 | FN50 | BioLegend | 1:50 |
| PD1 | BUV737 | EH12.1 | BD | 1:33 |
| FoxP3 | AF-647 | 206D | BioLegend | 1:50 |
| CD66b | PE | G10F5 | BioLegend | 1:40 |
| Siglec8 | PerCpCy5.5 | 7C9 | BioLegend | 1:20 |
| CD68 | FITC | Y1/82A | BioLegend | 1:12.5 |
| CD80 | BV711 | 2D10 | BioLegend | 1:50 |
| CD206 | APC | 15-2 | BioLegend | 1:20 |
| CD14 | BUV737 | M5E2 | BD | 1:20 |
| HLA-DR | BV510 | L234 | BioLegend | 1:20 |
| CD16 | BUV395 | 3G8 | BD | 1:100 |
| CD11c | BV605 | 3,9 | BioLegend | 1:20 |
| CD123 | BV421 | GH6 | BioLegend | 1:80 |
| PD-L1 | PE-Cy7 | B7-H1 | BioLegend | 1:20 |
| CD117 | BV785 | 104D2 | BioLegend | 1:20 |
| Neurophil phenotyping | | | | |
| CD45 | AF700 | HI30 | BioLegend | 1:100 |
| CD62L | BUV805 | DREG-56 | Termo fischer | 1:20 |
| CD163 | BV510 | GHI/61 | BioLegend | 1:20 |
| CD10 | BUV395 | HI10a | BD | 1:40 |
| CD63 | BV785 | H5C6 | BioLegend | 1:50 |
| CXCR4 | BV605 | 12G5 | BioLegend | 1:20 |
| CD11b | BUV737 | ICRF44 | Termo fischer | 1:50 |
| CD66b | BV421 | G10F5 | BD | 1:33 |
| Siglec8 | PerCpCy5 | 7C9 | BioLegend | 1:20 |
| CD107b | FITC | H4B4 | BioLegend | 1:50 |
| CD64 | BV711 | 10,1 | BioLegend | 1:50 |
| CD54 | PE/Dazzle | HA58 | BioLegend | 1:100 |
| Sialyl Lewis X | FITC | FH6 | BioLegend | 1:200 |
| CD274 | BV711 | 29E.2A3 | BioLegend | 1:100 |
| CD182 | PE/Dazzle | 5E8/CXCR2 | BioLegend | 1:200 |
| LDNs | | | | |
| Lox-1 | BV421 | 15C4 | Biolegend | 1:100 |
| CD10 | BUV395 | HI10a | BD | 1:40 |
| CD11b | BUV737 | ICRF44 | Termo fischer | 1:50 |
| PDL1 | BUV805 | MIH1 | BD | 1:50 |
| CXCR4 | BV605 | 12G5 | BioLegend | 1:20 |
| CD102 | FITC | CBR-IC2/2 | BioLegend | 1:100 |

**Table S2: List of antibodies used for IMC.**

| Mass Tag | Metal | Marker | Clone | Company | 5h/Overnight (ON) | dilution |
|---|---|---|---|---|---|---|
| 113 | In | CD31 | EP3095 | Abcam | 5h | 1:50 |
| 115 | In | pERK1/2 | D13.14.4E | CST | 5h | 1:100 |
| 140 | Ce | CD10 | MA5-14050 | Thermo fischer | 5h | 1:50 |
| 145 | Nd | pSTAT3 | M9C6 | CST | 5h | 1:50 |
| 146 | Nd | CD16 | EPR16784 | Abcam | 5h | 1:50 |
| 147 | Sm | CD163 | EDHu-1 | Novusbio | 5h | 1:100 |
| 152 | Sm | CD45 | D9M8I | CST | 5h | 1:50 |
| 154 | Sm | HIF1a | EP1215Y | Abcam | 5h | 1:50 |
| 156 | Gd | CD4 | EPR6855 | Abcam | 5h | 1:50 |
| 162 | Dy | CD8 | C8/144B | BioLegend | 5h | 1:100 |
| 166 | Er | CD263 | policlonal | ThermoFischer | 5h | 1:25 |
| 167 | Er | GranzymeB | EPR20129-217 | Abcam | 5h | 1:100 |
| 170 | Yb | CK7 | RCK105 | Abcam | 5h | 1:50 |
| 171 | Yb | CXCR4 | 12G5 | BioLegend | 5h | 1:100 |
| 172 | Yb | CD54 | HA58 | BioLegend | 5h | 1:50 |
| 139 | La | CD119 | polyclonal | ThermoFischer | ON | 1:50 |
| 141 | Pr | NE | 950302 | R&D | ON | 1:200 |
| 142 | Nd | CD63 | H5C6 | BD | ON | 1:200 |
| 143 | Nd | pP38 | 36/p38 | BD Biosciences | ON | 1:50 |
| 144 | Nd | pMAPKAPK2 | 27B7 | CST | ON | 1:100 |
| 148 | Nd | pS6 | D57.2.2E | CST | ON | 1:100 |
| 149 | Sm | CD191 | polyclonal | ThermoFischer | ON | 1:50 |
| 150 | Nd | pSTAT5 | C11C5 | CST | ON | 1:50 |
| 151 | Eu | PD-L1 | 28.8 | Abcam | ON | 1:100 |
| 153 | Eu | iNOS | SP126 | Abcam | ON | 1:100 |
| 155 | Gd | FOXP3 | 236A/E7 | Abcam | ON | 1:25 |
| 157 | Gd | CD87 | EPR2091 | Abcam | ON | 1:100 |
| 158 | Gd | CD33 | WM53 | BioLegend | ON | 1:100 |
| 159 | Tb | CitH3 | 7C10 | LSBio | ON | 1:50 |
| 160 | Gd | CD66b | BLR111H | Fluidigm | ON | 1:100 |
| 161 | Dy | MPO | E1E7I | SCT | ON | 1:200 |
| 163 | Dy | VEGF | G153-694 | BD pharmigen | ON | 1:100 |
| 164 | Dy | CD3 | D7A6E | CST | ON | 1:25 |
| 165 | Ho | pCREB | 87G3 | CST | ON | 1:50 |
| 168 | Er | Ki67 | B56 | BD | ON | 1:100 |
| 169 | Tm | MMP9 | L51/82 | BioLegend | ON | 1:200 |
| 173 | Yb | CD56 | MRQ-42 | Millipore Sigma | ON | 1:50 |
| 174 | Yb | HLADR | LN3 | BioLegend | ON | 1:200 |
| 175 | Lu | HLAG | 4H84 | BD | ON | 1:50 |
| 176 | Yb | HistoneH3 | D1H2 | Fluidigm | ON | 1:200 |
